# Alleviation of Drought Stress and Plant Growth Promotion in Mungbean through Rhizospheric Actinobacteria

**DOI:** 10.1101/2025.05.05.652240

**Authors:** Mrugesh M. Patel, N. K. Singh, Rudra B. Parmar, Bhavesh M. Joshi, Anurag Yadav, Yashvi R. Patel, Poonam V. Tapre

**Author notes:** Co-author.

## Abstract

Drought is a major cause for decreasing in crop yield and agricultural productivity around the Globe. The application of drought tolerance Actinobacteria to agroecosystem improves the growth and protect from water scarcity and they can be better alternative for natural farming and sustainable agriculture for aride and semi-aride regions. Herein, we isolated ten morphologically different actinobacteria from the rhizospheric soil of Datura and Khejri plants which are surviving in the nature without any special treatment provided. All the isolated bacteria were screened *in-vitro* for their drought tolerance capability for different time intervals against polyethylene glycol 8000 (PEG8000) with 5%, 10% and 10% concentration to media which produce moisture stress in which medium bacteria were growing. Among them best five bacterial isolates were selected and named RADM1 to RADM5 for further experimentation. Physiological characterizations were done through their ability to grow in different temperature, pH ranges and NaCl concentration supplemented with the 5% PEG8000. Morphology of the bacteria was Gram’s positive and mycelium producing same as typical Actinobacterial species and for biochemical diversity stains were able to IMVIC test and carbohydrate utilization through IMVIC test. Enzymatic observations of isolates were resulted positive for nitrate reduction, catalase production and negative for starch hydrolysis. PCR amplification and 16S *r*RNA gene sequencing in BlastN found maximum similarity of three species with *Streptomyces clavuligerus* and two species with *Rhodococcus erythropolis* and submitted to NCBI gene bank database with accession number PQ114139 (RADM1), PQ120343 (RADM2), PQ114109 (RADM3), PQ120390 (RADM4), PQ039766 (RADM5). In pot trial of mungbean, all the isolates showed better performance in bacterial treatments with an average increase in physiological parameters like, seedling’s shoot length (52.70% and 9.62%), root length (35.63% and 17.25%), fresh weight (42.8% and 13.76%) dry weight (45.25% and 21.24%) in compared to uninoculated induced drought and watered treatments. Stress features of seedlings revealed significant increase in proline and chlorophyll contents whereas decrease in MDA contents than the untreated plants which indicates actinobacteria reduced water scarcity and damage of plant membrane.

**IMPORTANCE:** Rhizosphere of plant contains diversity of microbes with different abilities to support the plant growth. The role of plant growth promoting bacteria in soil presented and their association with plant can be the one reason for survive of the plant against the abiotic stress conditions like drought, salinity, temperature or pH. The actinobacteria found in the rhizosphere have evolved multiple strategies to assist plants in managing the adverse effects of drought. These strategies may include alterations in phytohormones, which play a vital role in enabling plants to withstand environmental stresses, modifications to root structures, accumulation of osmolytes, enhancements in the plant’s antioxidant defense mechanisms, synthesis of exopolysaccharides, and the identification of particular genes that support plant growth and improve drought resilience. These actinobacteria as bioinoculant can be the better alternative for the chemical fertilizers use and protect from the water scarcity.

## INTRODUCTION

Drought stress is one of the major agricultural problems reducing crop yield in arid and semiarid regions of the world. As world’s climatic condition changes the mean global air temperature and precipitation patterns are leading to longer drought period and more extreme dry years and more severe drought conditions may hinder food production in some countries (Lau and Lennon, 2012). At present, strategies to increase the ability of plants to tolerate drought stress involve the use of water saving irrigation, traditional breeding and genetic engineering of drought tolerant transgenic plants. Unfortunately, these methods are highly technical and labour intensive and thus difficult to apply in practices.

To eradicate the problem of water scarcity, modern agro-biotechnological strategies are being implemented to improve drought tolerance in plants. These technologies include genetic engineering, germplasm screening, plant breeding, etc., which have resulted in the development of hybrid products that are widely grown across different parts of the Globe (Quan *et al.,* 2004). Nevertheless, abiotic stress tolerance mechanisms are complex, making the task of introducing new tolerant varieties tough and strenuous (Naveed, 2013).

Alternative agricultural strategies aimed at mitigating the adverse impacts of drought include conservation tillage, soil enhancement, and mulching. However, these approaches are often characterized by their complexity and the significant time investment they require (Bardi and Malusà, 2012). As agricultural practices extend into less fertile and marginal lands, there is a growing focus on interventions that enhance water use efficiency in crops, utilizing biotechnology and advanced agronomic techniques to meet the rising food demand.

Bacteria have developed various strategies to help plants cope with the damage caused by drought. These strategies can involve changes in phytohormones, which are crucial for helping plants deal with environmental stresses, modifications in the structure of plant roots, the buildup of osmolytes, adjustments in the plant’s antioxidant defense systems, the production of exopolysaccharides, and the identification of specific genes that promote plant growth and enhance drought resistance (Bechtold and Field, 2018).

Several studies have reported different solutions to the problematic situation of chemical usage in agriculture. Among the proposed solutions, microorganisms with multifunctional traits have been found to reduce the use of chemical fertilizers and pesticides by producing or releasing different types of bioactive compounds (Janardhan *et al.,* 2014), enzymes (Turan *et al.,* 2016), and antimicrobial substances or biocontrol compounds (Dhanasekaran *et al.,* 2005). In addition, others are plant growth promoters (Sarikhani *et al*., 2020). Considering this, plant growth-promoting rhizobacteria (PGPR) are a good alternative, and they have demonstrated mutualistic relationships with plants (Tang *et al*., 2016;).

The recent emphasis on utilizing beneficial microbial species that promote plant growth to mitigate the negative impacts of drought has gained significant attraction in agricultural practices. Bacteria play a crucial role in soil ecosystems, forming mutualistic and advantageous relationships with a majority of plant species (Ndeddy Aka and Babalola, 2017). These symbiotic bacteria enhance stress tolerance in various plant hosts through mechanisms such as phytohormones alterations, exopolysaccharide production, osmolytes accumulation, and defense against reactive oxygen species (Zhang *et al*., 2008). Furthermore, they possess the ability to synthesize antibiotics, fix atmospheric nitrogen, produce soluble iron compounds (siderophores), and solubilize inorganic phosphate (Babalola, 2010).

Plant growth-promoting bacteria (PGPB), a specific group of soil microorganisms, are instrumental in helping plants withstand the adverse effects of various ecological stresses (Vardharajula *et al*., 2011). These bacteria contribute to plant growth by directly supplying phytohormones, including gibberellins, cytokinins, and indole-acetic acid, while also facilitating the absorption of vital nutrients such as phosphorus and atmospheric nitrogen (Alori and Babalola, 2018). Furthermore, PGPB can serve as biocontrol agents, mitigating the damage caused by pathogenic fungi or bacteria. Therefore, it is imperative to explore and enhance plant growth and productivity under conditions of restricted water availability through the utilization of PGPB.

One alternative for growing plants under dry conditions is the use of plant growth promoting rhizobacteria (PGPR). PGPR are a group of bacteria that can be found in the rhizosphere in association with plan root system, both at the root surface and in endophytic associations and which can either directly or indirectly facilitate optimal plant growth and tolerance to biotic or abiotic stress condition (Cassan *et al*., 2009).

These outstanding properties of the plant growth promoting bacteria (PGPB) facilitate the plant growth during unfavourable environmental condition like drought (Yandigeri *et al*., 2012). The use of actinomycetes species to enhance stress tolerance in plant have receives very little attention over the years. Actinomycetes, found mostly in soils, are widely known for their antibiotic and bioactive secondary metabolites production as well as their outstanding ability to survive in unfavourable environments (Adegboye and Babalola, 2013).

Actinobacteria, a type of unicellular Gram-positive bacteria, are widely distributed in nature from different habitats and are important producers of several bioactive secondary metabolites, antibiotics, and plant growth-promoting factors (Pepper and Gentry, 2015) Actinobacteria aremorphologically very similar to fungi, though they form hyphae much smaller than fungi. The phylum Actinobacteria is considered one of the important groups of bacteria (Manivasagan *et al.,* 2014).

Today’s world requires ecofriendly methods to achieve high output yield, enhance crop production and better soil fertility (Yasari *et al*., 2009). Actinobacteria, as bio-inoculants and bio-pesticide, are an alternative to chemical fertilizers. They can improve crop production under multiple stress conditions, such as temperature, pH, salinity and drought (Cheng *et al*., 2018). *Streptomyces* is the most abundantly occurring actinobacteria genus in the soil. Owing to their high growth rate, *Streptomyces* spp. efficiently colonizes the plant root systems and can withstand adverse growth circumstances through the formation of spores.

Understanding the diversity and distribution of indigenous actinobacteria in the rhizosphere of particular crops depends on the knowledge of native Actinobacterial populations, their isolation, identification, and characterization. It is therefore mandatory to explore region specific Actinobacterial strains that can be used as growth promoters to achieve desired crop production (Deepa *et. al*., 2010). However, in spite of high soil population, secondary metabolite production and capability to endure hostile environments, *Streptomyces*, and other actinobacteria are unexpectedly under explored for plant-growth promotion, as compared to *Pseudomonas* or *Bacillus* spp. (Doumbou *et al.,* 2002).

Moreover, metagenomics studies have confirmed that actinobacteria have higher diversity in organic soil than in conventional modern agricultural soil (Sharma *et al*., 2019). Owing to the high price of chemical fertilizers and the widening gap between supply and demand, the solubilization of nutrients by the microorganisms has been increasingly seen to be useful and economical. Microbial inoculants are ecofriendly and environmentally secure and only require low cost technology, providing a solution for enhancing productivity and decreasing environmental problems for sustainable agriculture.

Soil Actinobacteria are known to produce active compounds in the rhizosphere, many of which are important in agriculture (Suzuki *et al*., 2000). Khan *et al*., (2010) reported that phosphorus could be implicated in several metabolic processes of plant hosts, such as energy transfer, photosynthesis, macromolecular biosynthesis, signal transduction, and respiration. Phosphorus availability to plants is facilitated through the soil P cycle. Richardson and Simpson (2011) noted that Actinobacteria directly solubilize and mineralize inorganic P or mediate organic P availability through microbial turnover and development of root system. Actinobacteria lower soil pH through the secretion of different types of organic acids, improving P availability to plants and in turn plant yield through the establishment and development of the entire root system shoots (Khan *et al*., 2010).

Here, we concentrate on actinobacteria as an alternative tool for sustainable farming practices and for reducing harmful chemical usage to promote ecofriendly sustainable agriculture, thus reducing environmental damage and transforming chemical farming into organic farming. The interaction of actinobacteria in relation to plant growth development, secondary metabolite production, biocontrol activity, and soil nutrient management.

In present study rhizospere of datura and khejri plant have been used as the source soil for the isolation of actinobacteria. The rhizospheric soil of these plants are selected for isolation of actinobacteria because these plants are of hardy nature and are found in abundance in wild in the North Gujarat Agroclimatic zone. These plants are further acclimatized to tolerate the dry and arid environmental conditions prevalent over the North Gujarat and are supposed to harbor beneficial actinobacteria in their rhizospheric zone and may have diverse plant growth promoting abilities.

## 1. Methods and Materials

### 1.1 Isolation of Actinobacteria from rhizospheric soil

#### 1.1.1 Collection of samples

The soil samples for the isolation of Actinobacteria were collected from rhizosphere of two plants growing wild in the North Gujarat region under arid condition; Datura (*Datura stramonium*) and Khejri (*Prosopsis cineraria*) from a depth of 15-30 cm (Datura) and 30-45 cm (Khejri). About 250g of soil from the each rhizosphere of Datura and Khejri’s were aseptically collected in sterile plastic bags. These soil samples were stored in refrigerator of the Department of Microbiology until further analysis.

#### 1.1.2 Isolation of Actinobacteria

All the soil samples collected from the different rhizospheres of datura and khejri were respectively combined and two separate composite samples of two plants were prepared. The samples were separately mixed thoroughly and passed through 2mm sieve to prepare homogenized samples. Ten grams of each soil samples were suspended in 90mL of sterile physiological saline solution (0.85% NaCl in distilled water) in a bottle and kept for shaking on an orbital shaker (at 100 rpm) at 28 ± 2 0C for 1h. After shaking, the samples were serially diluted up to 10-5 dilutions and aliquot from 10-4 and 10-5 dilutions (0.1mL) were spread plated on petriplates having Actinomycetes Isolation Agar (AIA) medium (Hi-Media Laboratories, Mumbai, India) supplemented with cycloheximide (20mg/l) and nalidixic acid (100mg/l) to minimize fungal and other bacterial growth. The petriplates were incubated at 28 ± 2°C for 3 to 7 days. Post incubation, all the plates were screened for actinomycetes colonies based on morphology and Gram staining. The Gram-positive bacteria showing typical morphological characters of actinobacteria (Cappucino and Sherman, 1992) on Actinomycetes Isolation Agar under microscope were isolated on freshly prepared actinomycetes agar plates. These cultures were further purified by repeated subculturing on International Streptomyces Project (ISP2) agar plates to obtain pure cultures. These pure cultures of actinobacteria were stored on ISP2 agar slants and ISP2 plates in refrigerators at 40C for further experimentation.

### 1.2 Screening of actinobacterial isolates based on drought tolerance capability

The drought tolerance capacity of the actinobacterial isolates were studied by inoculating 5 ml of freshly prepared actinobacterial aliquot into 150 ml cotton plugged flasks containing 50 ml sterilized ISP-1 medium (Tryptone Yeast Extract HiVegTM Broth) supplemented with different concentrations (5, 10 and 15%) of Poly Ethylene Glycol 8000 (PEG 8000) with slightly modification in method of Chukwuneme *et al*. (2020). The pH of the medium were adjusted to 7.2 before autoclaving at 121°C for 15 min. Inoculated flasks were incubated at room temperature with constant shaking (150 rpm) at different time intervals (24, 48 and 72 h). The growths of each bacterial isolates were observed by measuring the OD at 600 nm using a UV Spectrophotometer. The control for this experiment was consisted of inoculated broth without PEG8000, while un-inoculated broth served as blank. The pooled data were statistically analysed.

### 1.3 Physiological characterization of actinobacterial isolates

#### 1.3.1 Effect of temperature on the growth of bacteria

The bacterial ability to tolerate higher temperature were observed by growing 10µl of each bacterial isolate in 10 ml of sterilized ISP-1 medium containing 5% PEG-8000 and incubated at different temperatures (25°C, 30°C, 35°C, 40°C) under shaking conditions (150 rpm) for 7 days according to method described by Chukwuneme *et al*. (2020) with slightly modification. The OD_600_ of each bacterial isolate were measured at 600 nm using a UV Spectrophotometer. The OD observed was indicated the effect of temperature on growth of actinobacteria in medium with water stress.

#### 1.3.2 Effect of pH on the growth of bacteria

The effect of pH on bacterial growth were determined by growing 10 µl of each bacterial culture in test tubes containing 10 ml of sterilized ISP-1 medium supplemented with 5% PEG 8000. The pH of the media was adjusted to 3, 5, 7, 9, and 11 using 0.1 N HCl and 0.1 N NaOH before autoclaving. Inoculated tubes were incubated at 25°C under shaking conditions (150 rpm) for 7 days according to method described by Chukwuneme *et al*. (2020) with slightly modification.Aafter which the OD_600_ of each culture were measured using a UV spectrophotometer.

#### 1.3.3 Effect of NaCl on the growth of bacteria isolates

Tolerance to NaCl by bacteria were assessed according to the method of Ndeddy Aka and Babalola (2017) by inoculating 20 µl of each bacterial isolate in 20ml sterilized ISP-1 medium containing varying concentrations of NaCl (0.2, 0.4, 0.6, 0.8, and 1.0%). Inoculated tubes were incubated at 25°C for 7 days, and the OD were measured at 600 nm using a UV spectrophotometer according to method described by Chukwuneme *et al*. (2020) with slightly modification. An OD_600_ values greater than 0.1 was considered to be better growth for each bacterial isolates.

### 1.4 Morphological characterization

The microscopic examination of the isolates was done by Gram’s staining procedure and the observations for cell shape and arrangement of cells in colonies were recorded using the method given by Cappuccino and Sherman (1992) using a compound light microscope (E200, Nikon, Japan).

### 1.5 Biochemical characterization

#### 1.5.1 IMVIC test

Hi-Media ready to use IMVIC test kit is a standardized colorimetric identification system that utilizes four conventional biochemical tests and eight carbohydrate utilization tests. The tests are based on the principle of pH change and substrate utilization. On incubation organisms undergo metabolic changes which are indicated as a colour change in the media that can be either interpreted visually or after addition of the reagent. Indole test, methyl red test, Vogus-proskauer test, citrate utilization test, mannitol and sucrose test were carried out using IMVIC test kit (KB001-10KT, Hi-Media) following manufacturer protocol.

Single colony of a particular isolate was inoculated into 5 ml brain heart infusion broth and incubated at 35-37°C for 4 to 6 h. until the inoculum’s turbidity become ≥ 0.1 O.D. at 620 nm. Each well of the kit was inoculated with an aliquot of 50 μl of bacterial suspension and the plates were incubated at 35 - 37°C for 18 - 24 h. Reagents were added in well no. 1 (Kovac’s reagent), 2 (Methyl Red reagent) and 3 (Barrit reagent A and Barrit reagent B) at the end of incubation period, that is after 18-24 h. The result was observed in the form of colour change of the medium in wells of the kit as per manufacturer’s protocol.

#### 1.5.2 Nitrate reduction test

Bacterial suspension was inoculated into nitrate broth. Tubes were incubated at the optimal temperature 30°C or 37°C for 24 h. After incubation N2 gas was looked as bubble production before adding reagents. Then 6-8 drops of nitrite reagent A were added and then 6-8 drops of nitrite reagent B were added. The reaction (colour development) was then observed for a minute. If no colour develops then zinc powder was added to it. After addition of zinc powder, it was observed for at least 3 minutes for red colour to develop. Development of a cherry red coloration on addition of reagent A and B or absence of a red colour development on adding Zn powder indicates positive test. Development of red colour on addition of Zn powder indicates negative test.

#### 1.5.3 Catalase production test

Luria-Bertani (LB) agar plates were inoculated with each bacterial isolate. An un inoculated LB plates were used as the control. All plates were incubated at 30°C for 5 days. On the 5th day, a single colony from each inoculated plate were placed on a sterile glass slide and held at an angle while 2-3 drops of hydrogen peroxide (H2O2) was flooded over the growth of each culture on the glass slide. Catalase productions were indicated by the production of oxygen bubbles within one minute after the addition of H2O2.

#### 1.5.4 Test for hydrolysis of starch

A loopfull of each bacterial isolate were streaked on starch agar plates in triplicate; plates were incubated at 30°C for 8 days. On the 8th day, the iodine solution was added to each culture plate. After about a minute, excess iodine was carefully poured out from the culture plates. Control consisted of un-inoculated starch agar plates. Yellow zones around the colony in a dark blue medium was considered positive for starch hydrolysis, while the absence of a yellow zone was considered negative.

### 1.6 Molecular characterization of actinobacterial isolates

#### 1.6.1 Genomic DNA Extraction

Bacterial isolates were grown in 20 ml of ISP-1 medium in 50 ml Eppendorf tubes with constant agitation (150 rpm) at a temperature of 25°C for 7 days for optimum growth. The genomic DNA from each bacterium was extracted using a DNA extraction kit (Qiagen) according to the manufacturer’s instructions. DNA purity and yield (ng/µl) were determined using a NanoDrop spectrophotometer.

#### 1.6.2 PCR amplification

The 16S *r*RNA gene is considered molecular chronometer due to the highly conserved nature of these genes. PCR amplification of the 16S *r*RNA gene was done using the universal primers 27F and 1492R in order to get an amplified PCR product of size ∼1500 bp using A300 thermal cycler (Long Gene Scientific Instruments Co. Ltd., Taiwan).

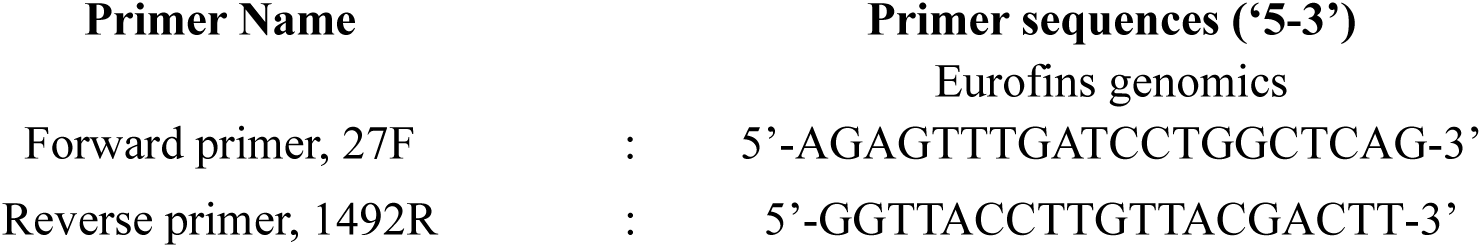

The reaction mixture was given a short spin in a microcenrifuge for mixing of the cocktail components. The PCR tubes were then loaded in a thermal cycler.

❖ Thermal cycling conditions for PCR amplification
| Sr. No. | Steps | Temperature (°C) | Duration |
| --- | --- | --- | --- |
| 1. | Initial-denaturation | : 94 | : 05 min |
| 2. | Denaturation | : 94 | : 30 sec |
| 3. | Annealing | : 54.5 | : 01min 30sec |
| 4. | Extension | : 72 | : 02 min |
| Repeated 35 times to step 2 |  |  |  |
| 5. | Final Extension | : 72 | : 10 min |
| 6. | Hold | : 4 | : - |

#### 1.6.3 Gene sequencing and identification of 16S *r*RNA gene

The analysis of sequences and construction of the phylogenetic tree were performed according to the methods described by Aremu and Babalola, 2015. The chromatograms obtained from the sequencing reaction were analysed for good quality sequences and were edited with Bio Edit Sequence Alignment Editor, and consensus sequences were generated. Blast search were done on the consensus sequences obtained in the NCBI database (www.ncbi.nlm.nih.gov) using the Basic Alignment Search Tool (BlastN) for homology for identification of the bacteria. Sequences obtained were deposited in GenBank. The 16S *r*RNA gene sequence based dendrogram were also prepared online using Neighbour Joining Method to depict the phylogenetic position of these isolates.

### 1.7 Pot Experiment

Actinobacterial isolates after being characterized based on phenotype and PGPR attributes were tested on mungbean under pot studies in order to find out the best isolate for promoting plant growth. The sandy loam soil was collected from the field. The soil was then be steam sterilized and plastic pots were disinfected with 4 per cent formaldehyde (formalin 40 EC) solution used for conducting experiments in pots.

Seven Treatments comprised with soil inoculation of actinobacterial isolates for mungbean for 21 days of experiment. Five seeds were sown to the each pot with five replications and transferred into growth chamber at 25°C with a photoperiod of 16/8h light/dark. After one week, the seedlings were inoculated with the actinobacterial suspension in sterilized tap water respectively with an inoculum load of 10^8^ CFU/ml. One week after bacterial inoculation water supply was terminated to produce drought in the pot for further seven days and then after drought treatment water irrigation resumed for one day, and plant health were photographed and assessed. For comparison one treatment was maintained as control *i.e*., in this case neither soil treatment with bioinoculant nor water supplies were given and another control treatment was continued with water supply during experiment without bacterial inoculation. After twenty two days of experiment plants with different treatments were harvested for biomass and length measurements. Leaves from each treatment were respectively collected and used for the measurement of free proline, MDA and chlorophyll content (Chen et al, 2017). The data obtained were analysed with MSTAT-C software and DNMRT (Duncan’s New Multiple Range Test).

## 2. Results

### 2.1 Isolation of Actinobacteria from rhizospheric soil of datura and khejri plants

The rhizospheric soil of datura and khejri were collected from a depth of 15-30 cm (datura) and 30-45 cm (khejri). Five distinct bacterial colonies from each of the two soil samples were isolated in pure culture through repeated subculturing from soil sample and were stored in refrigerator at 4°C for further experimentation. These isolates were marked sequentially 1 to 10. According to method described by Anwar et al 2016; one gram of soil sample mixed with 9ml of autoclaved water and was serially diluted to a final dilution of 10^−4^ and 10^−5^. The 0.1ml of each dilution was spread plated on Actinomycetes isolation agar medium supplemented with 25µl/ml nalidixic acid and 50µl/ml nystatin as antifungal agents. These ten actinobacterial isolates were selected on the basis of morphological characters such as distinct colour and colony shape. Morphologically distinct bacterial colonies that grew on the Actinomycetes isolation agar medium were aseptically transferred onto fresh ISP2 agar plates. These isolates underwent purification through repeated subculturing, The pure cultures were subsequently maintained on ISP2 agar slants.

### 2.2 Screening of actinobacterial isolates based on drought tolerance capacity and physiological characteristion

#### 2.2.1. Screening of actinobacterial isolates based on drought tolerance

The screening of the selected bacterial isolates to tolerate drought was evaluated based on concentration of PEG8000 and time. The level of tolerance to various concentrations of PEG8000 by each bacterial isolate was determined as a function of time, after inoculation on PEG containing medium. Growth varied among the isolates and depended mainly on the concentration of PEG, as bacterial growth decreased in PEG supplemented medium for all the isolates, irrespective of the concentration.

It was also observed that time influenced the tolerance capacities of these isolates as better growth on PEG medium was observed with an increase in time. The result obtained for each concentration was compared with that of the control (without PEG). Screening test were done after the isolation, growth of actinobacterial isolates against the drought tolerant capability measured with different concentration 5%, 10% and 15% of PEG8000 after 24h, 48h and 72h time intervals. Optical Density(OD) of growth of isolates in medium revealed that time influenced the tolerance (Table 5).

**Table 1:** Complete Randomized Design with 5%, 10% and 15% PEG concentration for 24h, 48h and 72h of incubation.

| Particulars | Values |
| --- | --- |
| Treatment | 10 levels |
| Hours | 3 levels |
| Total observations | 90 |

**Table 2:** Combined CRD analysis with 5% PEG8000 after 24h, 48h and 72h of incubation.

| Source of Var. | DF | Sum of Sq. | Mean Sq. | F value | P value | Result |
| --- | --- | --- | --- | --- | --- | --- |
| Treatment | 9 | 0.5509 | 0.0612 | 306.0 | 0.0 | ** Sig. at 1% level |
| Hours | 2 | 0.9735 | 0.4868 | 2434.0 | 0.0 | ** Sig. at 1% level |
| TreatmentxHours | 18 | 0.133 | 0.0074 | 37.0 | 0.0 | ** Sig. at 1% level |
| Error | 60 | 0.0133 | 0.0002 |  |  |  |
| Total | 89 | 1.6707 |  |  |  |  |

The means of all the levels of the interaction between treatments and hours were same.

The treatment component p=value was 0.0, results were found ** Sig. at 1% level and reject the null hypothesis, so multiple comparison test must be performed to evaluate means of different levels of the interaction between Treatment & Hours.

The interaction of isolates with 5% PEG after 24h, 48h and 72h of incubation revealed that isolates found more tolerance for drought after 72h after incubation than 24h and 48h incubation periods. Among all the interaction isolate T9 had maximum tolerance against PEG8000 after 72 h of incubation (O.D.= 0.8473±0.0025) and none of the treatments were at par with it. It was followed by isolates T8, T10, T4 and T2 with growth 0.793±0.0026, 0.765±0.004, 0.758±0.0017, 0.7453±0.0021 (Table 5).

**Table 3:** Combined CRD analysis with 10% PEG8000 after 24h, 48h and 72h of incubation.

| Source of Var. | DF | Sum of Sq. | Mean Sq. | F value | P value | Result |
| --- | --- | --- | --- | --- | --- | --- |
| Treatment | 9 | 0.6389 | 0.071 | 142.0 | 0.0 | ** Sig. at 1% level |
| Hours | 2 | 1.0064 | 0.5032 | 1006.4 | 0.0 | ** Sig. at 1% level |
| TreatmentxHours | 18 | 0.1412 | 0.0078 | 15.6 | 0.0 | ** Sig. at 1% level |
| Error | 60 | 0.0314 | 0.0005 |  |  |  |
| Total | 89 | 1.8179 |  |  |  |  |

The means of all the levels of the interaction between treatments and hours were same.

The interaction of isolates with 10% PEG after 24h, 48h and 72h of incubation revealed that different isolates shows different level of tolerance for growth after 24h, 48h and 72h of incubation ranges from 0.538±0.0031 to 0.014±0.003. Among all the interaction isolate T10 found highest tolerance against PEG after 72h (0.538±0.0031) and none of the treatments were at par with it, which were followed by isolate T9 with 48h (0.472±0.0006) and 72h (0.470±0.0032) of incubation. Least tolerance was observed in isolate T3 after 24h of incubation (0.014±0.003) followed by isolate T6 after 24h of incubation (0.021±0.0061) (Table 5).

**Table 4:** Combined CRD analysis with 15% PEG8000 after 24h, 48h and 72h of incubation.

| Source of Var. | DF | Sum of Sq. | Mean Sq. | F value | P value | Result |
| --- | --- | --- | --- | --- | --- | --- |
| Treatment | 9 | 0.6929 | 0.077 | 770.0 | 0.0 | ** Sig. at 1% level |
| Hours | 2 | 0.3092 | 0.1546 | 1546 | 0.0 | ** Sig. at 1% level |
| TreatmentxHours | 18 | 0.1229 | 0.0068 | 68.0 | 0.0 | ** Sig. at 1% level |
| Error | 60 | 0.0078 | 0.0001 |  |  |  |
| Total | 89 | 1.1328 |  |  |  |  |

The means of all the levels of the interaction between treatments and hours were same

**Table 5:**
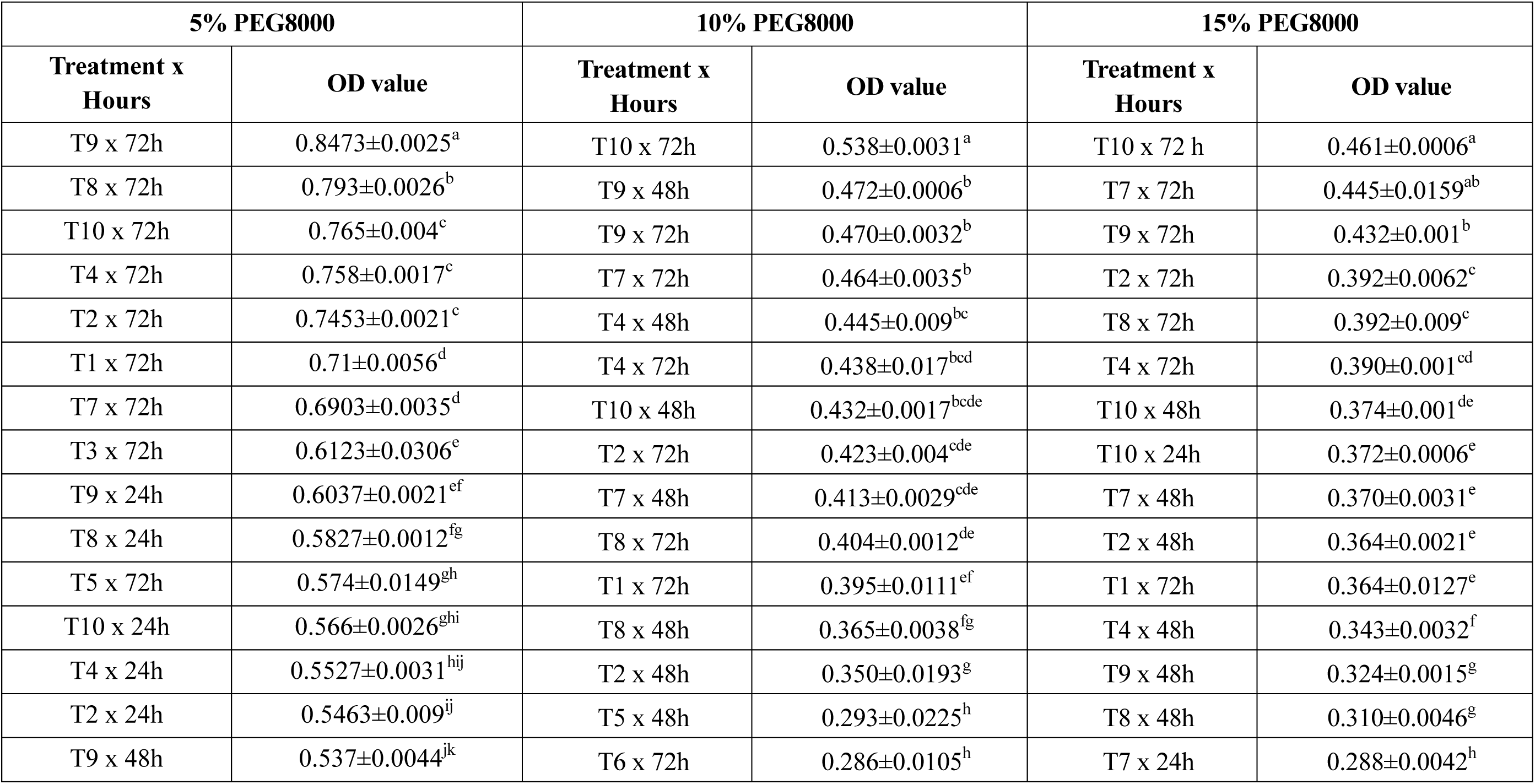
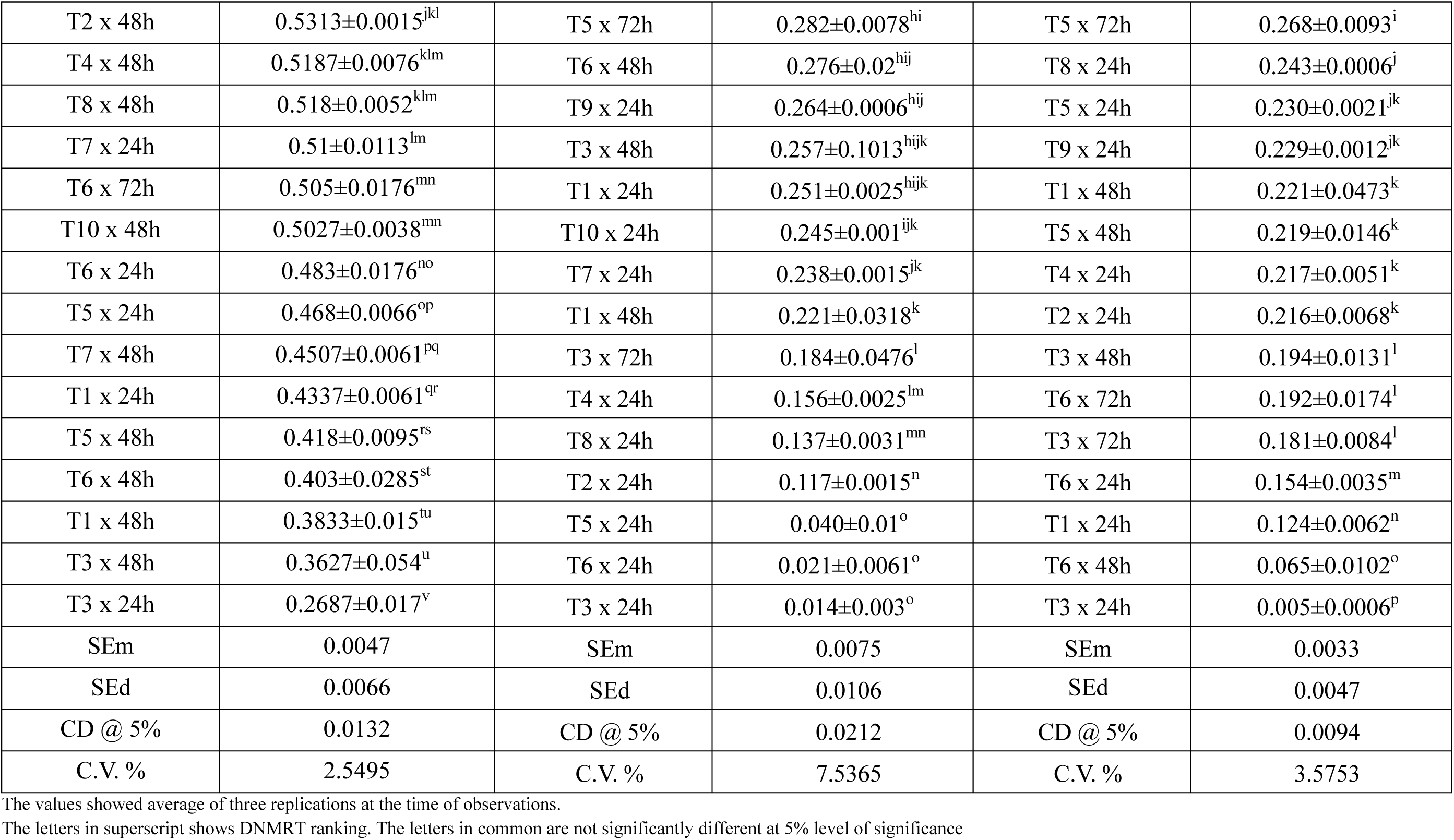
DNMRT test for comparing means of interactions of actinobacterial isolates against 422 5%, 10% and 15% PEG8000 for 24h, 48h 423 and 72h of incubation.

Comparison of tolerance for the growth of isolates at different concentration after different time of interval revealed that the least concentration of PEG8000 at 5% shows better growth than 10% and 15% concentration in medium. Isolate T9 had maximum tolerance to 5% PEG (0.06627) and isolate T3 had least tolerance to 15% PEG (0.127) (Table 6).

**Table 6:**
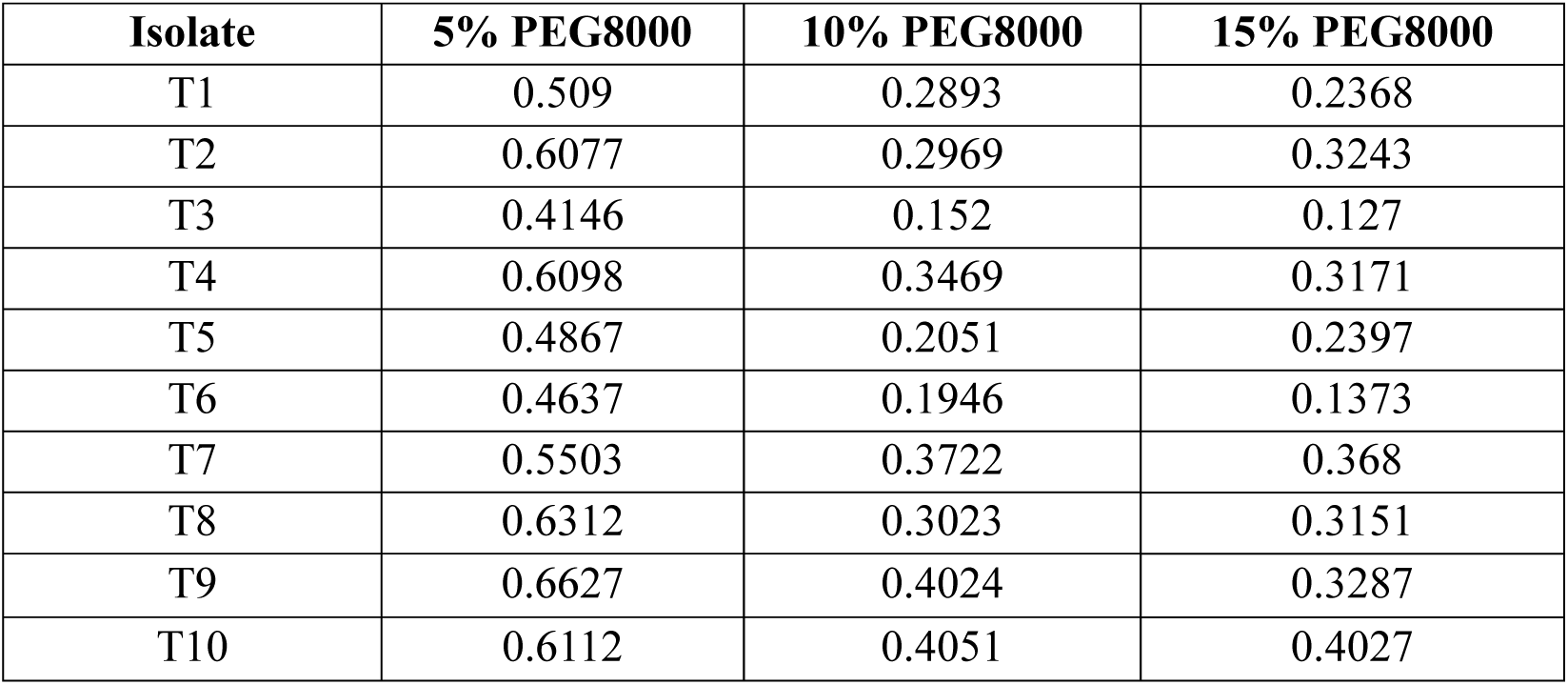
Mean OD value of isolates at 5%, 10% and 15% PEG8000 concentration after 24h, 48h and 72h after incubation.

| Isolate | 5% PEG8000 | 10% PEG8000 | 15% PEG8000 |
| --- | --- | --- | --- |
| T1 | 0.509 | 0.2893 | 0.2368 |
| T2 | 0.6077 | 0.2969 | 0.3243 |
| T3 | 0.4146 | 0.152 | 0.127 |
| T4 | 0.6098 | 0.3469 | 0.3171 |
| T5 | 0.4867 | 0.2051 | 0.2397 |
| T6 | 0.4637 | 0.1946 | 0.1373 |
| T7 | 0.5503 | 0.3722 | 0.368 |
| T8 | 0.6312 | 0.3023 | 0.3151 |
| T9 | 0.6627 | 0.4024 | 0.3287 |
| T10 | 0.6112 | 0.4051 | 0.4027 |

So that, the five Actinobacterial isolates which showed maximum tolerance against PEG8000 at 5% concentration according to the DNMRT value of comparing mean with different time interaction (Table 5) were selected and named as RADM1 [T9×72h (0.8473±0.0025 at 72h)], RADM2 [T8×72h (0.8473±0.0025 at 72h)], RADM3 [T10×72h (0.765±0.004 at 72h)], RADM4 [T4×72h (0.758±0.0017at 72h)] and RADM5 [T2×72h (0.7453±0.0021 at 72h)] for the further experiments.

### 2.2 Physiological characterization of actinobacterial isolates

#### 2.2.1 Effect of temperature on the growth of bacteria

The activity and growth of bacteria in the soil depend on the soil temperature, which affects cellular enzymes. In present study, five actinobacterial isolates after screening for drought tolerance were grown under medium supplemented with 5% PEG8000 and at different temperature regimes. The results obtained for each of the isolates at various temperatures are presented in Table 7 Bacterial growth was determined by measuring the optical density (OD) of each bacterium at 600 nm. All tested bacterial isolates grew at various temperatures, although there were variations in the growth pattern of each isolate.

**Table 7:**
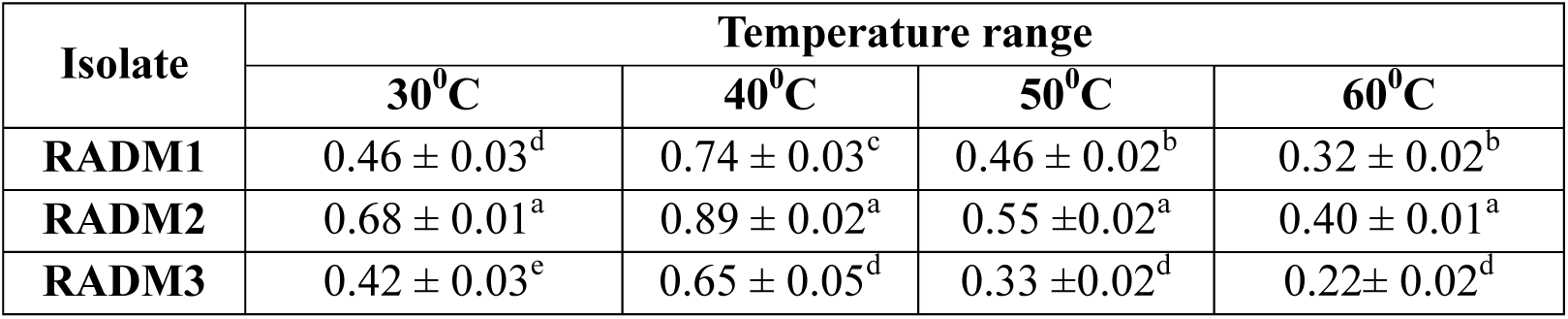
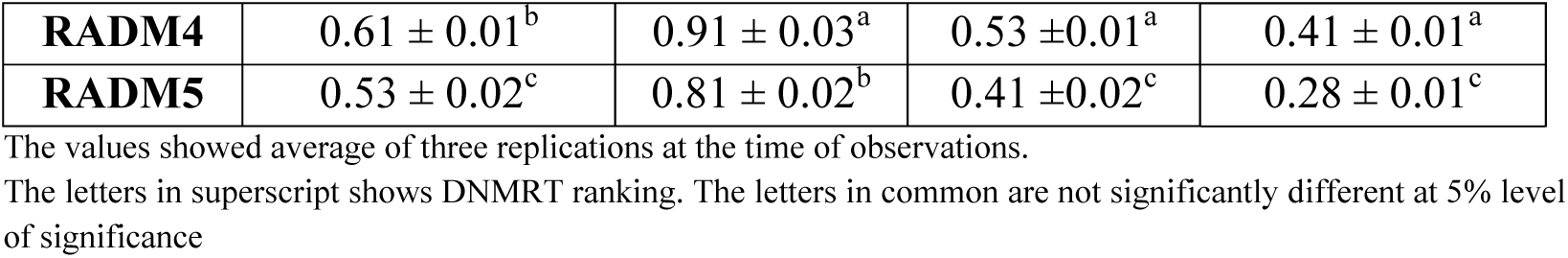
Growth of bacterial isolates at different temperature.

All the isolates showed maximum growth at 40°C on compared to growth at 30°C, 50°C and 60°C. Among them, isolate RADM4 has highest growth (0.91 ± 0.03) which was followed by the isolate RADM2 (0.89 ± 0.025) and the least growth was observed in isolate RADM5 (0.81 ± 0.025). At incubation temperature of 60°C, a significant decrease in the growth was observed for all the isolates. As per the result was observed by the Ndeddy aka and Babalola (2017) decreased the growth of bacterial isolates was observed when temperature decreased to 30° C or less and raise to 50° C or more. According to them, the decrease in growth at this temperature could be due to reduction in metabolic activity of bacterial isolates caused by the low/high temperature.

#### 2.2.2 Effect of pH on the growth of bacteria

The level of microbial activity in the soil is usually affected by the pH of soil. In the present study actinobacterial strains were grown under medium supplemented with 5% PEG8000 and at different pH ranges of 3, 5, 7, 9, 11 (Table 8). The result revealed that optimum growth at OD_600_ for all tested bacterial isolates was observed between the pH of 5and 9, All the actinobacterial isolates shows highest growth At 7 pH than other pH. pH of the environment influence bacterial survival and growth. For most soil bacteria, the specific pH range is usually between 4 and 9, with the optimum being 6.5 to 7.5 (Akond et al., 2016).

**Table 8:**
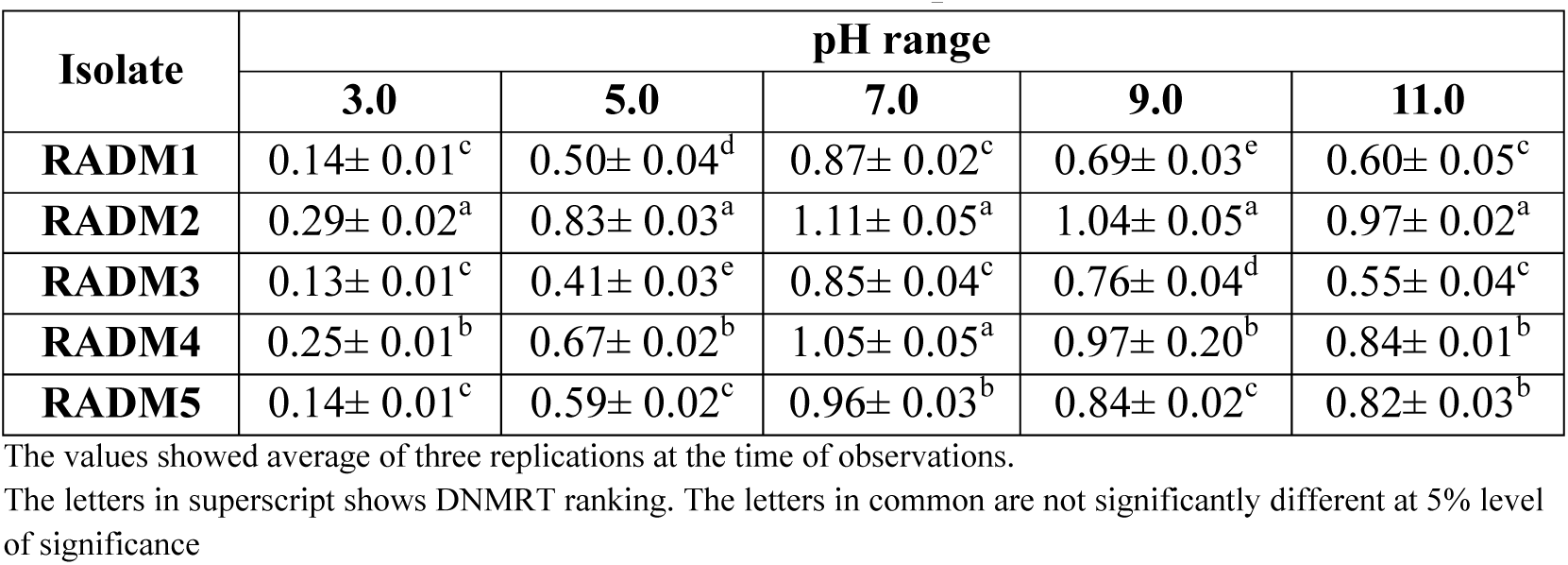
Growth of bacterial isolates at different pH.

Among them, the isolate RADM2 (1.11± 0.05) and RADM4 (1.05± 0.05) showed significantly higher growth compared to other isolates at pH 7 and the least growth was observed in the isolate RADM3 (0.85± 0.04). Isolate RADM2 was able to withstand better than the other isolates at all the pH under investigated. Even at pH 11 and pH3, the isolate RADM2 performed best and regestred OD value 0.97± 0.025 and 0.29± 0.02, respectively which were significantly higher than the other isolates at these pH values.

#### 2.2.3 Effect of NaCl concentration on the growth of bacteria

All the bacterial isolates were assessed for their ability to withstand salinity stress by growing at different concentration of sodium chloride (NaCl) 0.5, 1.0, 2.0, 3.0, 5.0 with the media supplemented with 5% PEG (Table 9). Significant differences were observed in the growth of bacterial isolates at different concentration of NaCl. At 0.5% NaCl the isolate RADM2 regestred highest growth (OD= 0.96± 0.04) followed by the isolate RADM4 (OD= 0.95± 0.043) and RADM1 (OD= 0.88± 0.025), respectively.

**Table 9:**
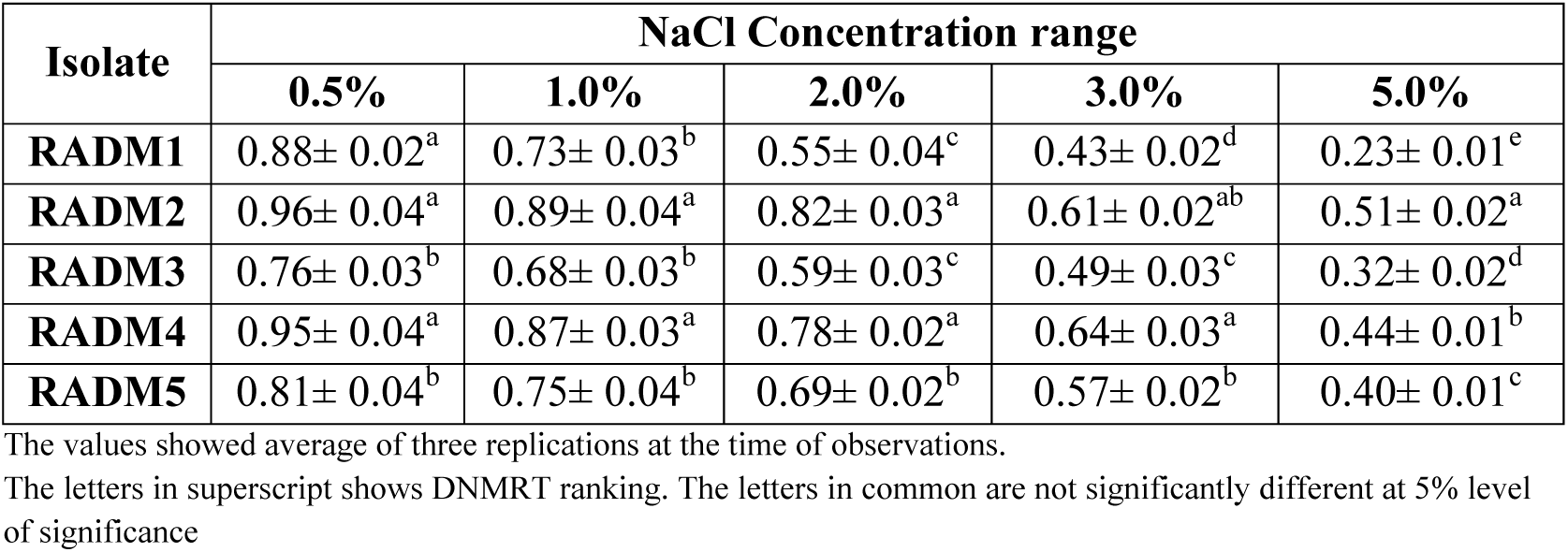
Growth of bacterial isolates at different NaCl concentration.

Growth of actinobacterial isolates respectively decreased with the increase in concentration of NaCl which could be an indication of its halotolerant nature. The least growth was observed for all the isolates at 5.0% NaCl concentration in media supplemented with 5% PEG8000 which optimised with both salinity and water stress.

### 2.3 Morphological characterization of isolates

Morphology has been an important characteristic to identify Actinobacteria isolates. This was done using various standard culture media, including International Streptomyces Project (ISP) medium. Various morphological observations, including germination of spores, elongation and branching of vegetative mycelium, formation of aerial growth, colour of colony and pigment production have been used to identify actinobacteria.

Microscopic examination of the isolates was done by Gram’s staining using the methodologies of Cappuccino and Sherman (1992). All the actinobacterial isolates were found Gram’s positive and purple coloured which is a distinct characteristic of Actinobacteria. These actinobacterial isolates were examined for their cell shapes under microscope. Among them, isolates RADM1, RADM2 and RADM4 were found rod shaped and isolates RADM3 and RADM5 were found to possess both variable rod and coccus shaped cells after 48h of growth on petriplate having ISP2 media ((Figure_1).

**Figure 1:**
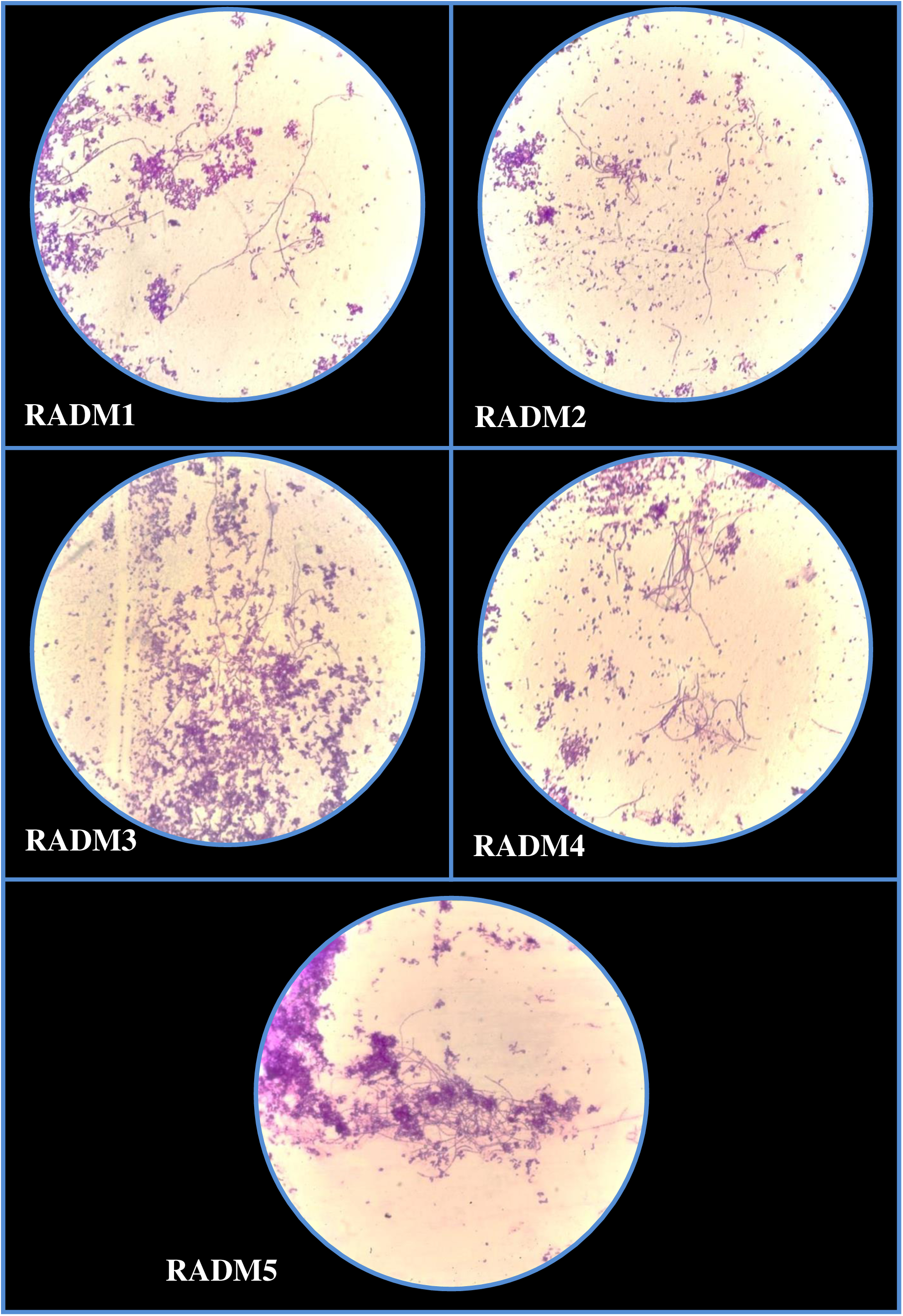
Microscopic examination of mycelium of actinobacterial isolates

Classic actinobacteria have well developed radial mycelium. According to the difference of morphology and functions, the actinobacterial mycelia can be divided into substrate mycelium and aerial mycelium. The substrate mycelia can be white, yellow, orange, red, green, blue, purple, brown or black; some hyphae can produce water soluble or fat soluble pigments. Aerial hyphae of actinobacteria depend upon the species characteristics, nutritional conditions or environmental factors.

Mycelium of Isolate RADM1, RADM2 and RADM4 were found filamentous, branched or unbranched and clumps at the end of hyphae. Branched hyphae were observed in the isolates RADM3 and RADM5 under microscopic examination after 120h of growth on ISP2 culture media ((Figure_1).

Cultural characteristics of isolates were observed on the petriplates prepared with ISP2 (Yeast extract malt extract agar) medium. Actinobacterial isolates appeared compact, often leathery, giving a conical appearance with dry surface on culture media. Pigmentation of actinobacteria varied (red, black, purple, white, grey, orange, yellow) according to species characteristics, nutritional conditions or environmental factors. Yellow to orange colour with dry surface were observed on petriplates streaked with actinobacterial isolates ((Figure_2). The strains produced yellowish orange to yellowish brown or olive substrate mycelium on ISP2 culture media. All the isolates were observed on irregular form and filamentous margins on cultured petriplates as indented peripheral edges on culture growth (Figure_2).

**Figure 2:**
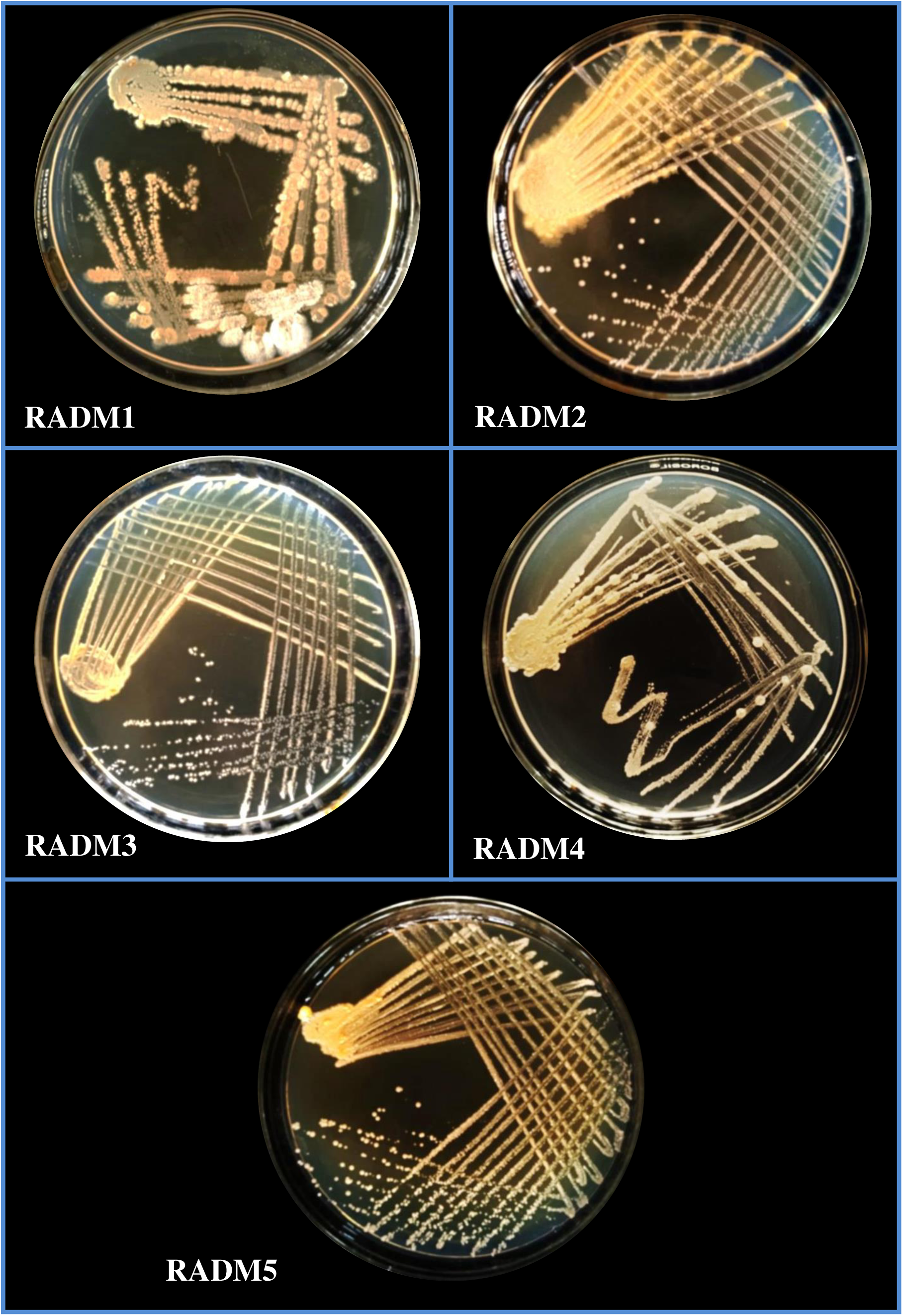
Pure culture of actinobacterial isolates on ISP2 medium

**Table 10:** Microscopic observation of actinobacteria isolates.

| Isolates | Gram's Staining | Cell Shape | Mycelial morphology |
| --- | --- | --- | --- |
| RADM1 | +Ve | Rod | Clumps, branched and unbranched hyphae |
| RADM2 | +Ve | Rod | Clumps, branched and unbranched hyphae |
| RADM3 | +Ve | Rod and coccus | Branched hyphae |
| RADM4 | +Ve | Rod | Clumps, branched and unbranched hyphae |
| RADM5 | +Ve | Rod and coccus | Branched hyphae |

**Table 11:**
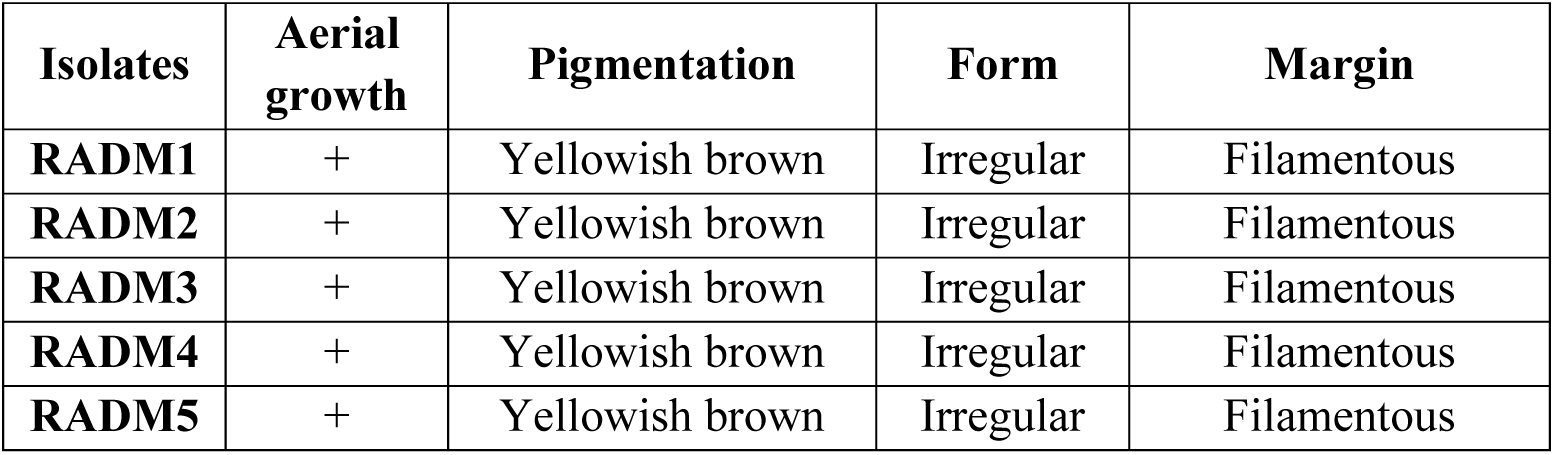
Cultural characterization of the actinobacteria.

### 2.4 Biochemical characterization of isolates

#### 2.4.1 IMVIC tests and carbohydrate utilization tests

Observations revealed that all the bacterial isolates were found positive for methyl red and glucose utilization and negative for indole, voges proskauer’s and sorbitol utilization. Isolate RADM1 and RADM2 found positive for citrate utilization arabinose, mannitol and sucrose utilization and negative for adonitol and lactose utilization. Isolate RADM3 were found positive for citrate and sucrose utilization, negative for mannitol and rhamnose utilization and produce intermediate reaction for adonitol, arbinose and lactose utilization. Isolate RADM4 were found positive for citrate, arbinose and mannitol utilization and negative for adonitol, lactose, rhamnose and sucrose utilization. Isolate RADM5 were found positive for methyl red and glucose utilization and negative for adonitol, arabinose, lactose, mannitol, rhamnose and sucrose utilization ((Figure_3, Table 12).

**Figure 3:**
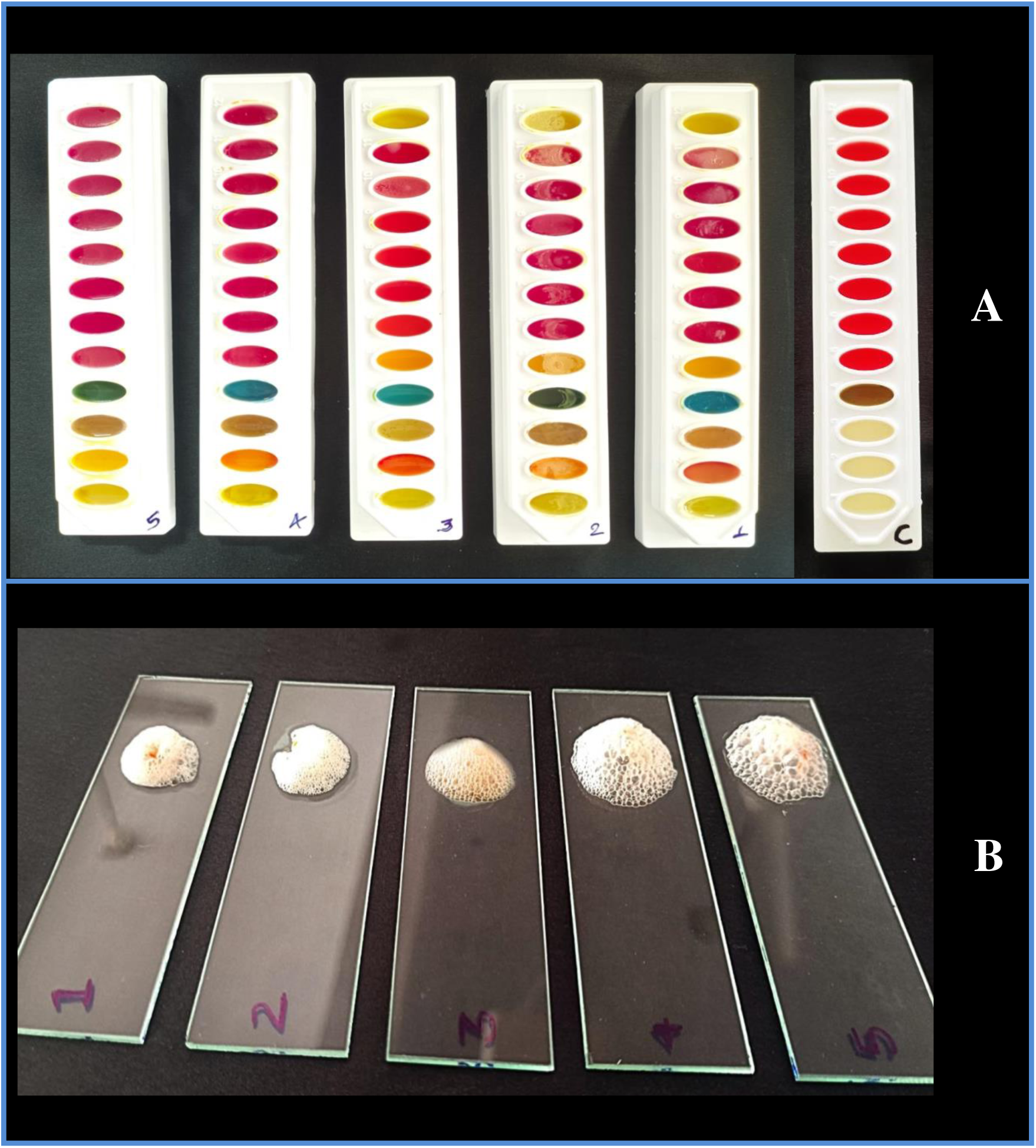
A- Biochemical characterization of actinobacterial isolates through IMVIC test kit B- *In-vitro* catalase test of actinobacterial isolates

**Table 12:** Biochemical characterization of Actinobacteria isolates.

| Sr.no. | Test | RADM1 | RADM2 | RADM3 | RADM4 | RADM5 |
| --- | --- | --- | --- | --- | --- | --- |
| **IMVIC tests** |  |  |  |  |  |  |
| 1 | Indole | - | - | - | - | - |
| 2 | Methyl red | + | + | + | + | + |
| 3 | Voges Proskauer's | - | - | - | - | - |
| 4 | Citrate utilization | + | - | + | + | - |
| **Carbohydrate Utilization tests** |  |  |  |  |  |  |
| 5 | Glucose | + | + | + | + | + |
| 6 | Adonitol | - | - | ± | - | - |
| 7 | Arabinose | + | + | ± | + | - |
| 8 | Lactose | - | - | ± | - | - |
| 9 | Sorbitol | - | - | - | - | - |
| 10 | Mannitol | + | + | - | + | - |
| 11 | Rhamnose | ± | - | - | - | - |
| 12 | Sucrose | + | + | + | - | - |
| **Enzymatic tests** |  |  |  |  |  |  |
| 1 | Nitrate reduction | + | + | + | + | + |
| 2 | Starch hydrolysis | - | - | - | - | - |
| 3 | Catalase | + | + | + | + | + |

The actinobacterial isolates were characterized for their biochemical properties based on colorimetric observation involving IMVIC test and carbohydrate utilization tests were carried out using Hi-Media IMVIC test kits. The isolate which showed positive reaction for biochemical test and for utilization of carbohydrate was indicated by ‘+’, the isolate which showed negative reaction was indicated by ‘-’, and the isolate which showed intermediate reaction are indicated by ‘±’.

#### 2.4.2 Catalase test

This biochemical differentiates the bacteria into aerobic and anaerobic and indicated whether they are Gram positive or Gram negative. Bacterial species which are capable to producing catalase are also able to breakdown the hydrogen peroxide into oxygen and water, the oxygen gas resulting into formation of bubbles, indicating a positive result. All the actinobacterial isolates were found positive for catalase productions as they found to produce bubbles ((Figure_3) while adding hydrogen peroxide and proved that the bacterial isolates were Gram’s positive. The catalase enzyme helps bacteria detoxify by neutralizing the harmful effects of hydrogen peroxide. Hydrogen peroxide can cause oxidative damage to organisms, including DNA, lipids, and proteins.

#### 2.4.3 Nitrate reduction test

The nitrate reduction test determines if nitrate was reduced by the bacteria in the sample. The test uses reagent to produce a red colour if nitrate is present in the medium. If the medium doesn’t turn red, it could be because the nitrate wasn’t reduced. All the actinobacterial isolates were found positive for the nitrate reduction as they produced cherry red colour after addition of reagent A & B ((Figure_4).

**Figure 4:**
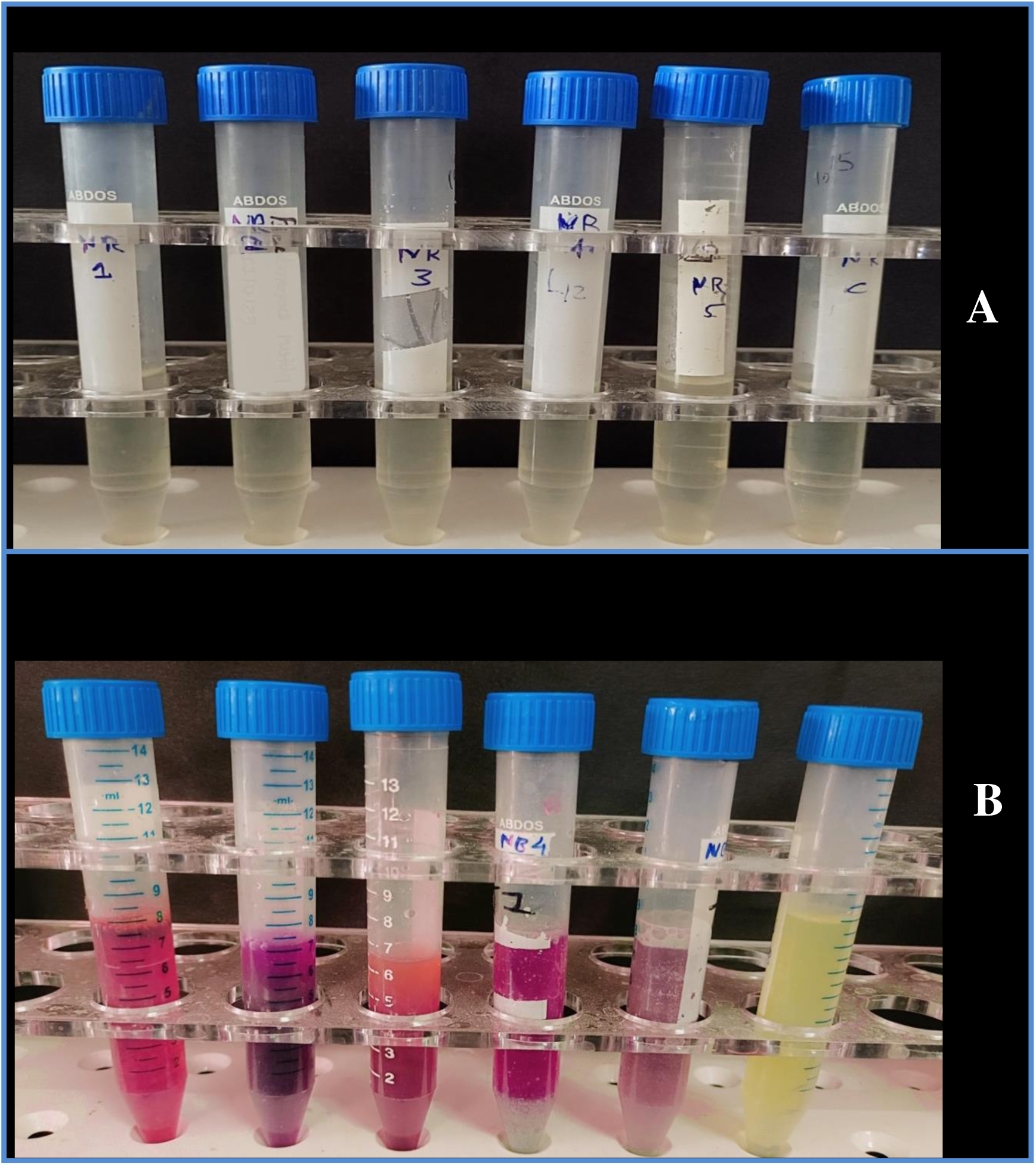
In-vitro nitrate reduction by actinobacterial isolates (A - before incubation, B – after incubation and addition of reagent A & B)

#### 2.4.4 Starch hydrolysis test

Bacteria that produce amylase secret this enzyme into the surrounding medium, where it breaks down the starch into smaller sugar molecules present in the medium, which results into production of a clear zone around bacterial colonies on starch agar plate after addition of iodine. Indicating clear halo around colonies where starch has been hydrolysed and is no longer present to react with iodine with uniform blue colour across the plate. All the bacterial isolates were recorded negative for starch hydrolysis test as they were unable to hydrolyse the starch molecules present in the media and no halo zone were observed after addition of iodine solution to petriplates ((Figure_5).

**Figure 5:**
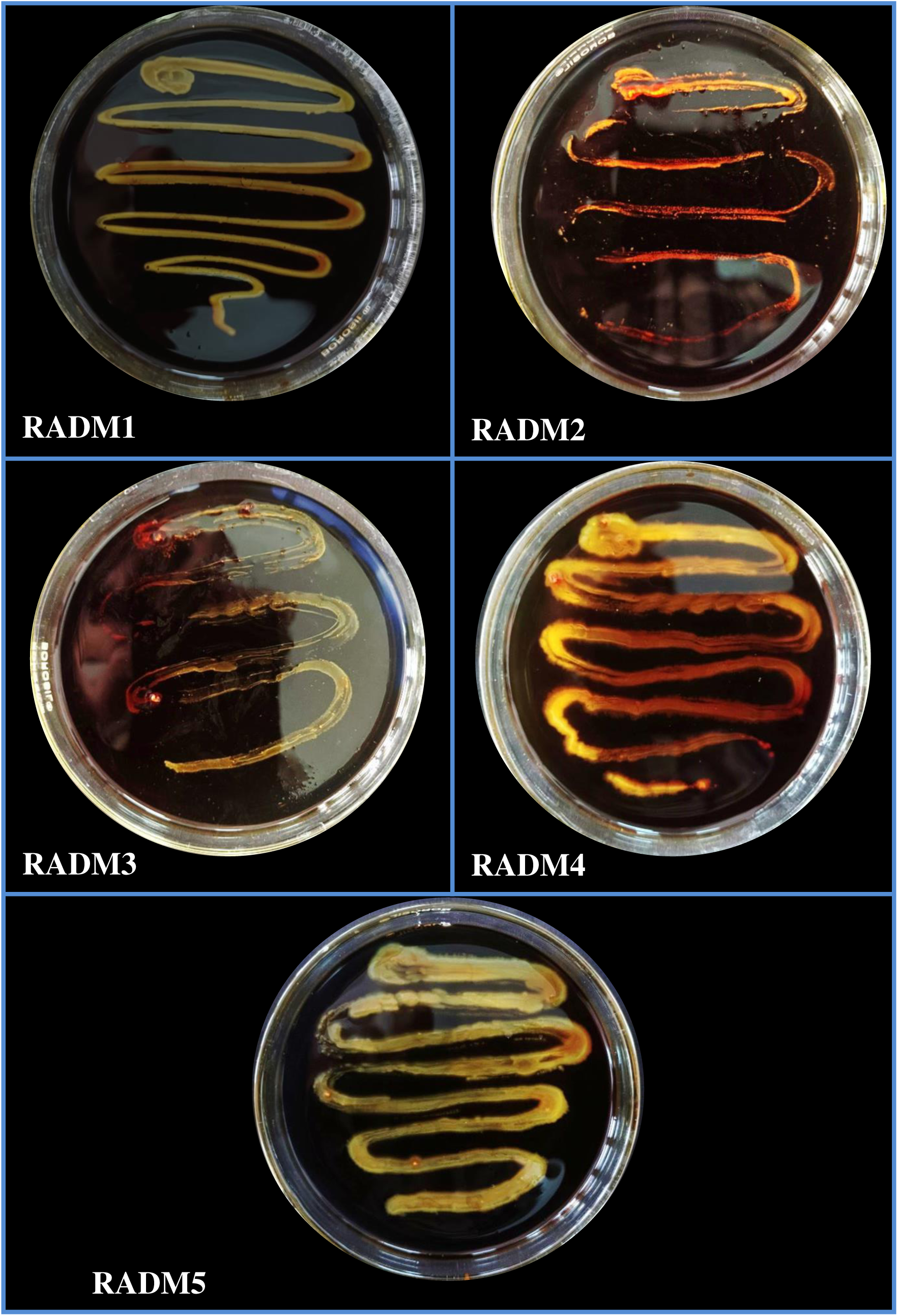
In-vitro starch hydrolysis by actinobacterial isolates

### 2.5 Molecular characterization of the actinobacterial isolates

Molecular characterization of bacteria plays a crucial role in accurately identifying bacterial species, determining their genetic relationship, tracking the spread of specific strains within a population and understanding their virulence factors, which is particularly important in different field of Microbiology. Unlike traditional phenotypic methods that can be ambiguous molecular techniques like DNA sequencing provide a highly accurate means to identify bacterial species even when they appear similar in morphologically.

#### 2.5.1 Spectrophotometric study of genomic DNA

The spectrophotometric analysis of DNA showed the absorbance ratio of DNA A_260_/A_280_ ranged from 1.63 to 1.86 with a mean value of 1.72. The DNA concentration ranged from 52.5 to 132.5, the average concentration of genomic DNA of the five isolates of actinobacteria was 84.67ng/µl. The entire five isolated DNA were treated with RNAase and proteinase K as per requirements to remove the nuisance in pure genomic DNA. Detailed information on quantification of DNA has been provided. Quality check of isolated genomic DNA was done by electrophoresing DNA samples in 0.8 per cent agarose gel stained with ethidium bromide (10 mg/ml) ((Figure_6). These DNA sample were further utilized for the polymerase chain reaction and identification of actinobacterial isolates obtained from the rhizosphere of datura and khejri plants through 16S *r*RNA gene amplification.

**Figure 6:**
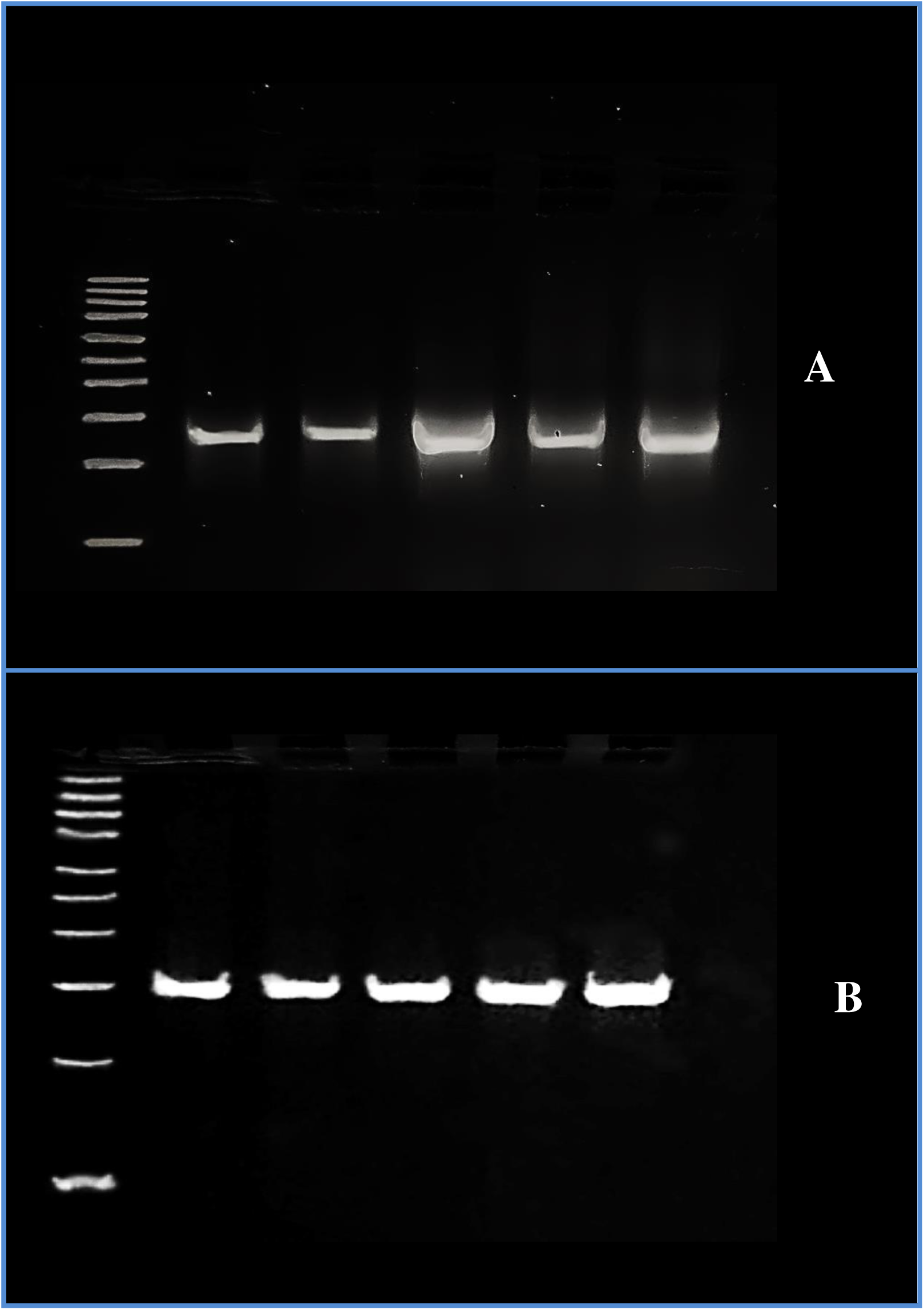
A: Genomic DNA of actinobacterial isolates B: PCR amplification of 16S *r*RNA gene with 27F and 1492R primer of actinobacterial isolates

#### 2.5.2 Polymerase chain reaction of the 16S *r*DNA gene

The 16S *rRNA* gene amplification by polymerase chain reaction with the 27F (5’- AGAGTTTGATCCTGGCTCAG-3’) and 1492R (5’-GGTTACCTTGTTACGACTT - 3’) primers revealed a band size of about 1.5Kb on 1.5% agarose gel stained with ethidium bromide in all the bacterial isolates ((Figure_6).

PCR amplification for 16S *r*RNA gene was also carried out by Bind *et al*. (2019) in 25 μL reaction volume having 2 μl DNA, 12.5 μl PCR master mix, 1.5 μl Forward primer, 1.5 μl reverse primer and 12.5 μl nuclease free water. Reaction mixtures were put to polymerase chain reaction (PCR) at optimized temperature conditions as: initial denaturation of the template DNA at 94°C for 5 min, denaturation at 94°C for 30 sec, annealing at 54.5°C for 1 min 30 sec, elongation at 72°C for 2 min and a final elongation at 72°C for 10 min. Denaturation, annealing and elongation cycles were repeated for 35 cycles. Amplification products were separated on a 1.5% (w/v) agarose gel in 1X TAE buffer and visualized by ethidium bromide staining. Amplification product of size about 1500bp using primers 27F and 1492R ((Figure_6).

#### 2.5.3 Gene sequencing and identification of the actinobacterial isolates

Identification of actinobacterial isolates were done by 16S *r*RNA gene sequencing and analysis. These isolates were sequenced using forward and reverse primers. The sequences obtained after alignment were compared using the BlastN tool, with the nucleotide sequences on NCBI GenBank database. The 16S *r*RNA gene sequences of five isolates were submitted to the NCBI GenBank and obtained accession numbers for submitted sequences (Table 13). Isolates RADM1, RADM2 and RADM4 were found respectively, 96.58%, 98% and 100% similarity with actinobacterial species *Streptomyces clavuligerus* of order Streptomycetales of phylum Actinomycetota. Isolates RADM3 and RADM5 were found respectively 98% and 96% similarity with the species of *Rhodococcus erythropolis* of order Mycobacteriales of phylum Actinomycetota (Table 13). These DNA sequences were submitted to NCBI were used to construct a phylogenetic tree with bootstrap values to decipher their evolutionary relationship among similar species based upon similarity and differences in their genetic characteristics ((Figure_7, 8, 9, 10, 11). The combined phylogenetic groupings of the actinobacterial isolates were prepared through Clustal Omega tool. ((Figure_12).

**Figure 7:**
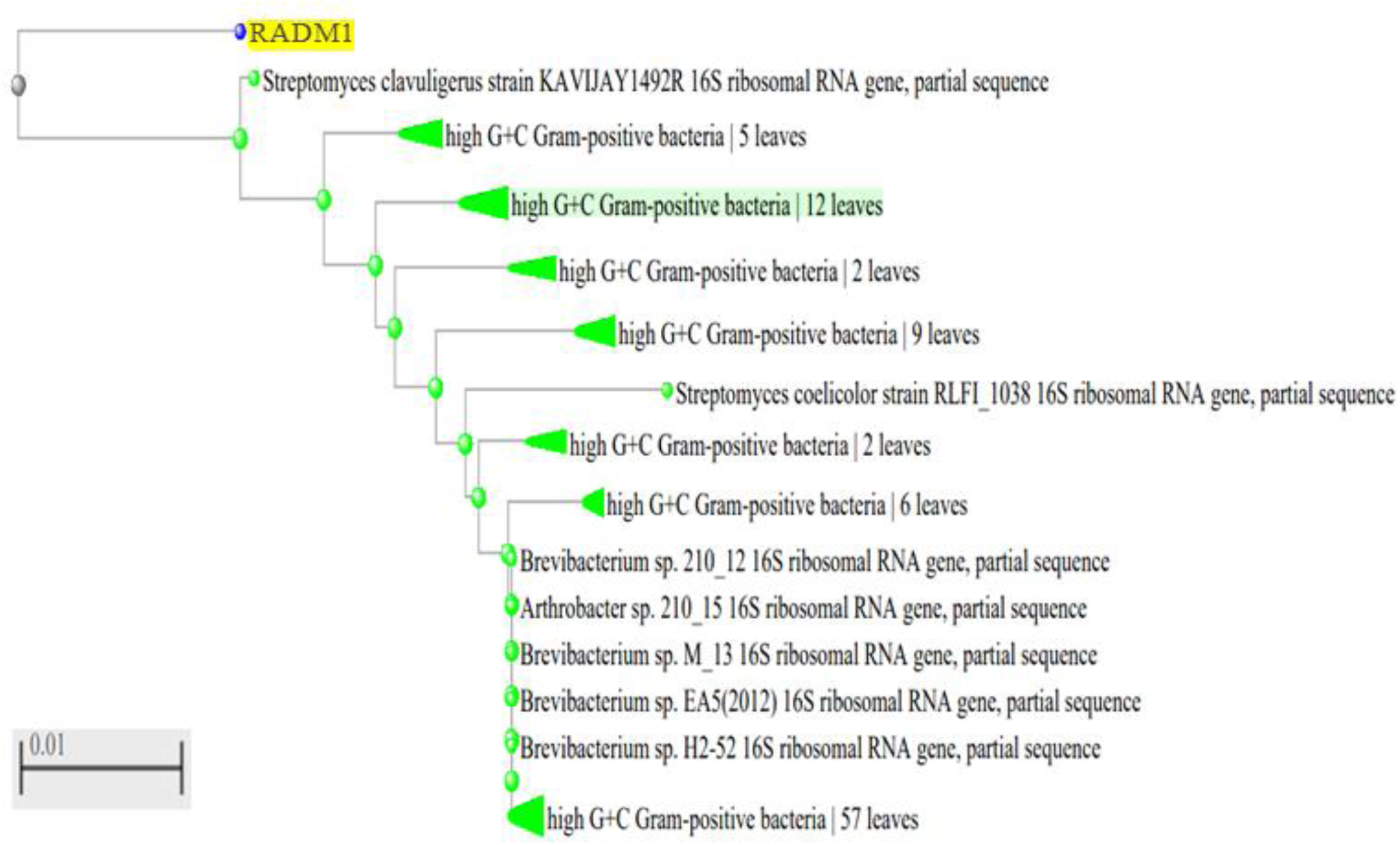
Phylogenetic grouping of isolate RADM1 by Neighbour joining method

**Figure 8:**
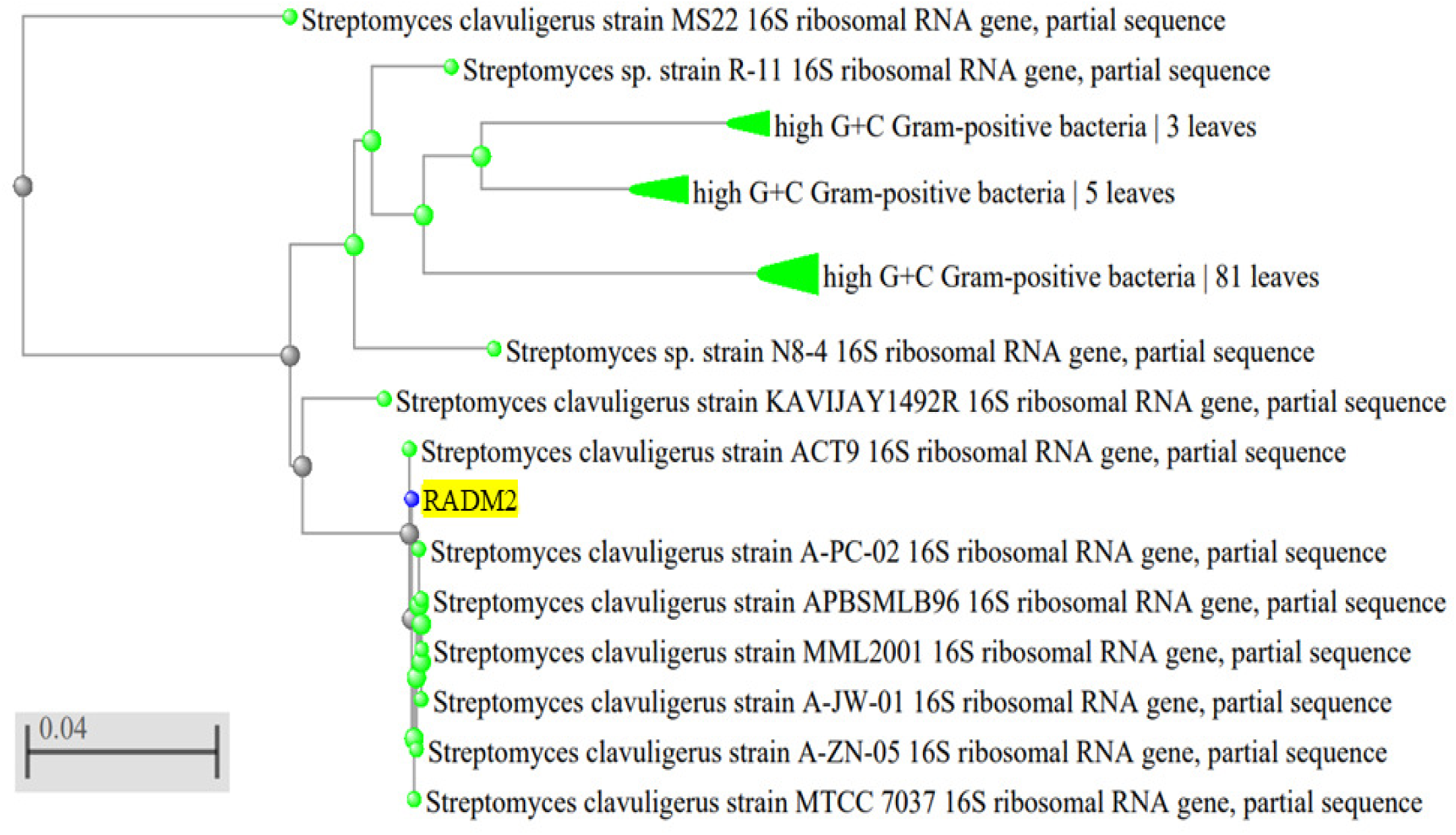
Phylogenetic grouping of isolate RADM2 by Neighbour joining method

**Figure 9:**
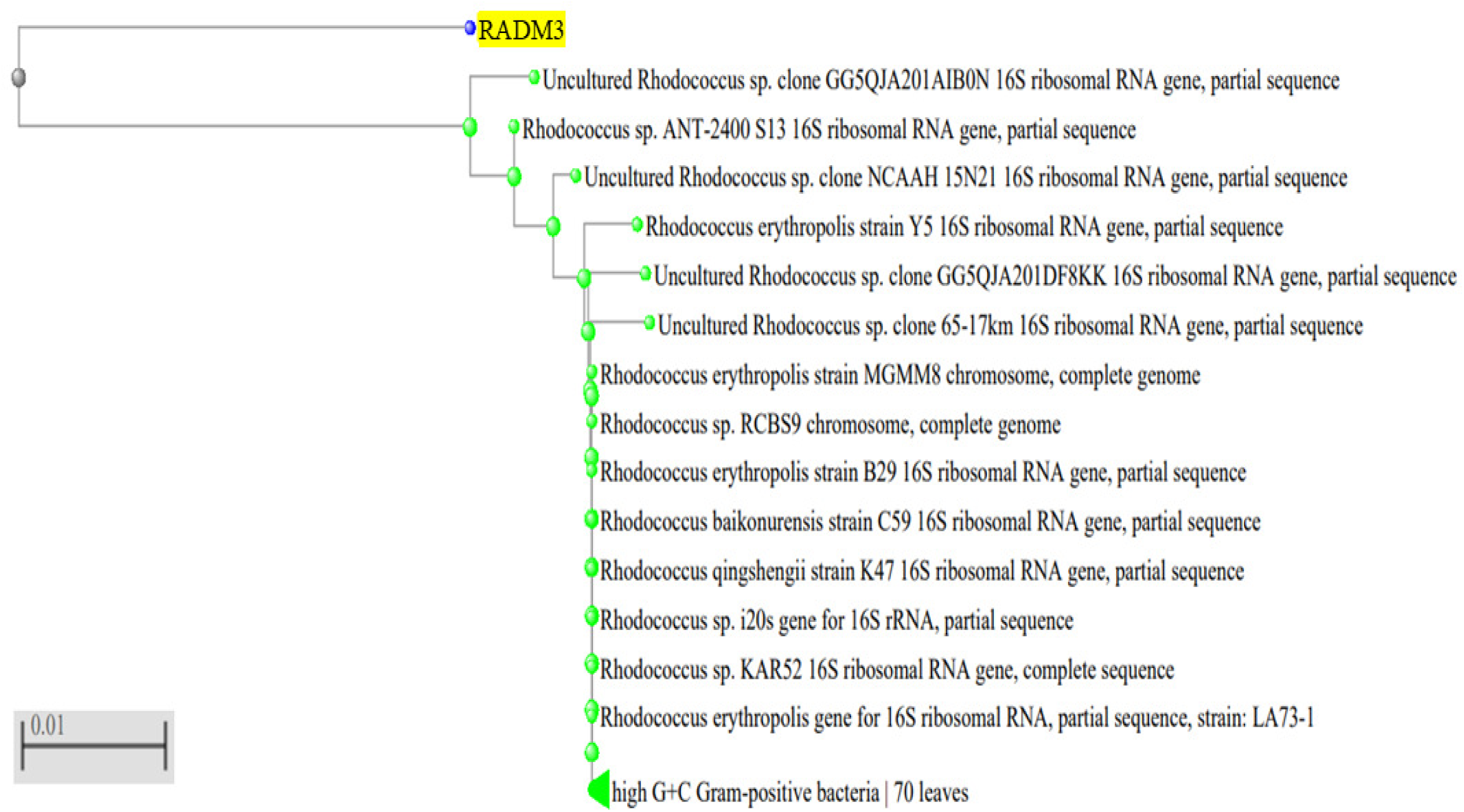
Phylogenetic grouping of isolate RADM3 by Neighbour joining method

**Figure 10:**
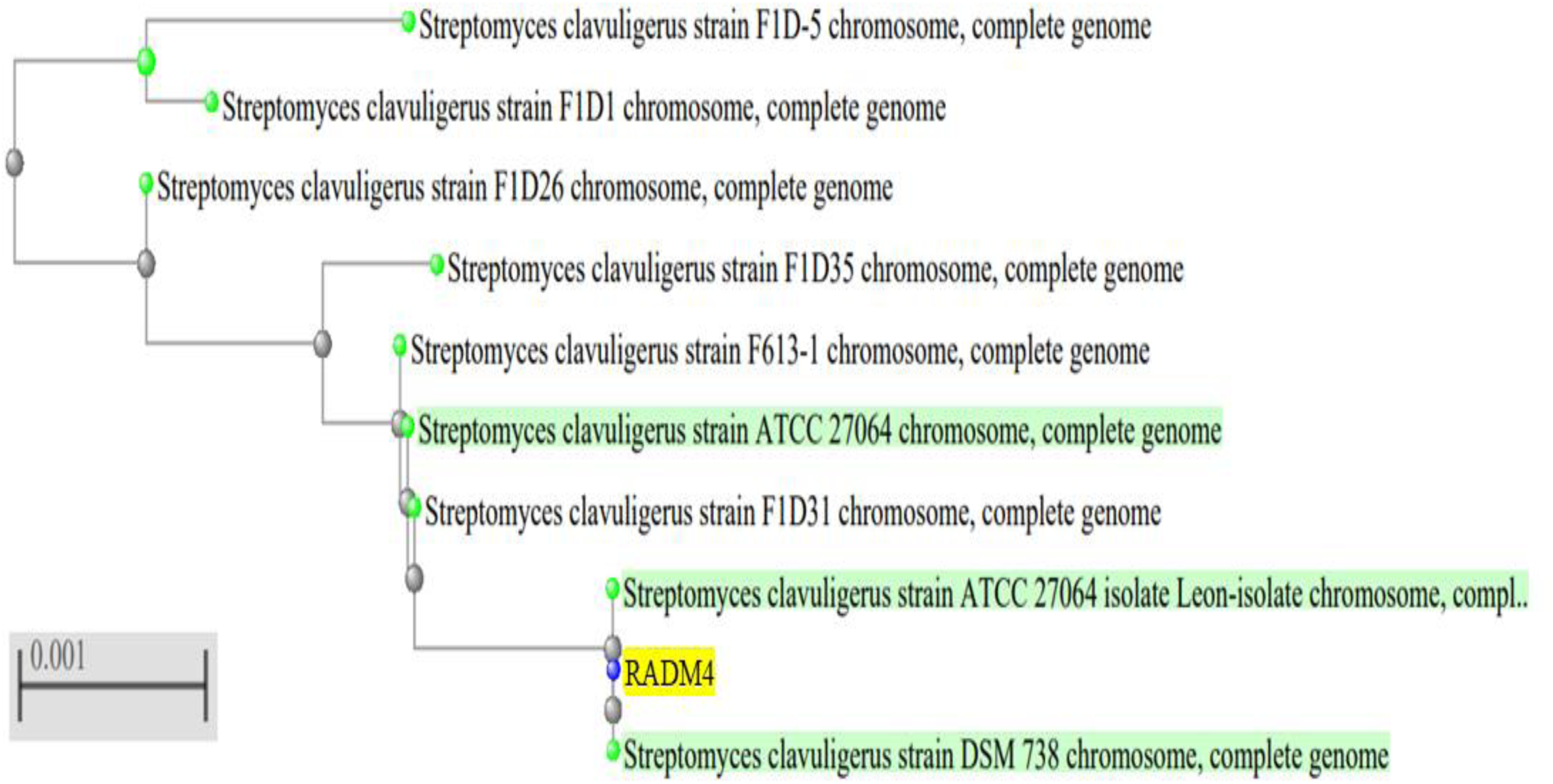
Phylogenetic grouping of isolate RADM4 by Neighbour joining method

**Figure 11:**
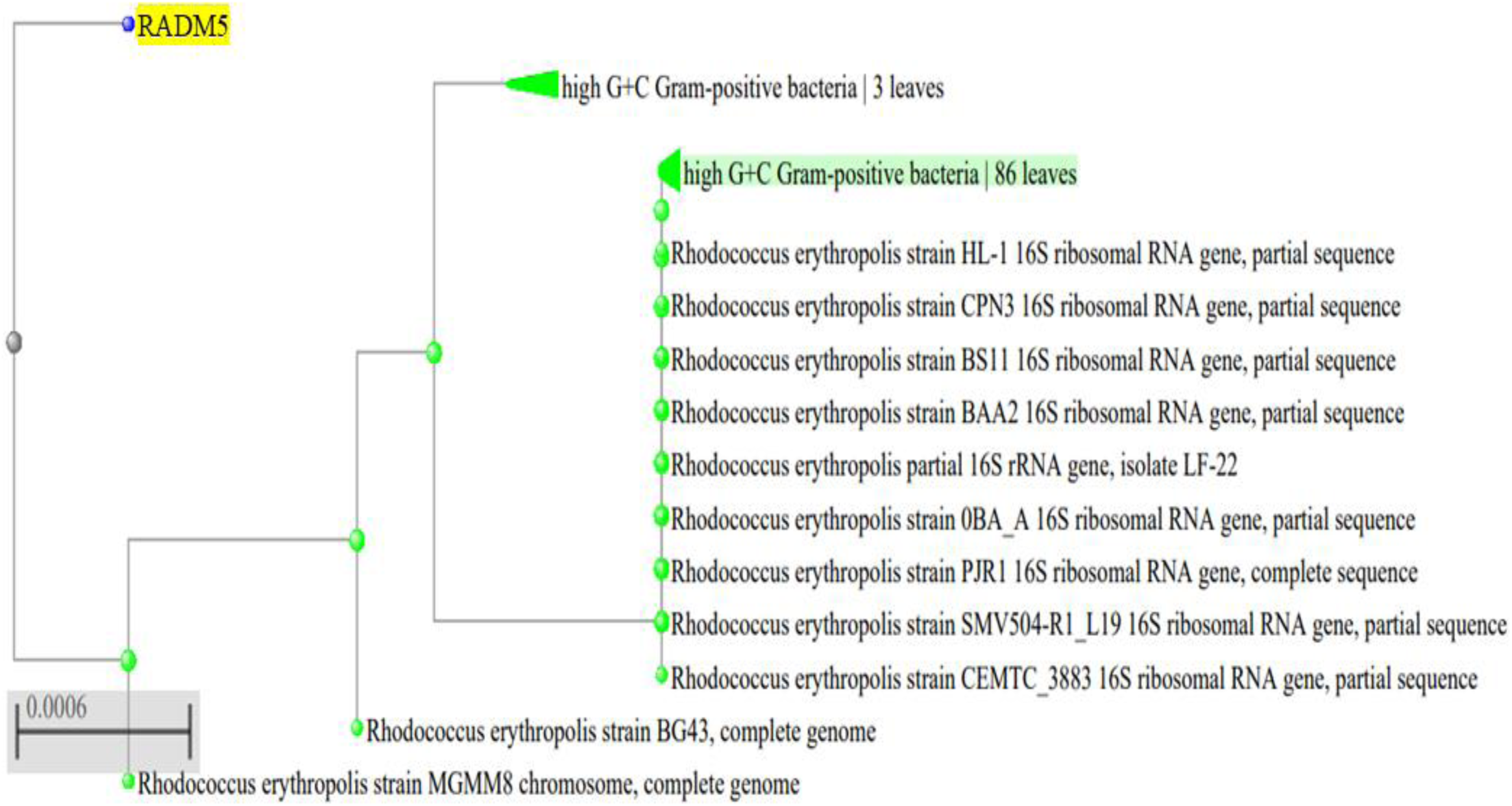
Phylogenetic grouping of isolate RADM5 by Neighbour joining method

**Figure 12:**
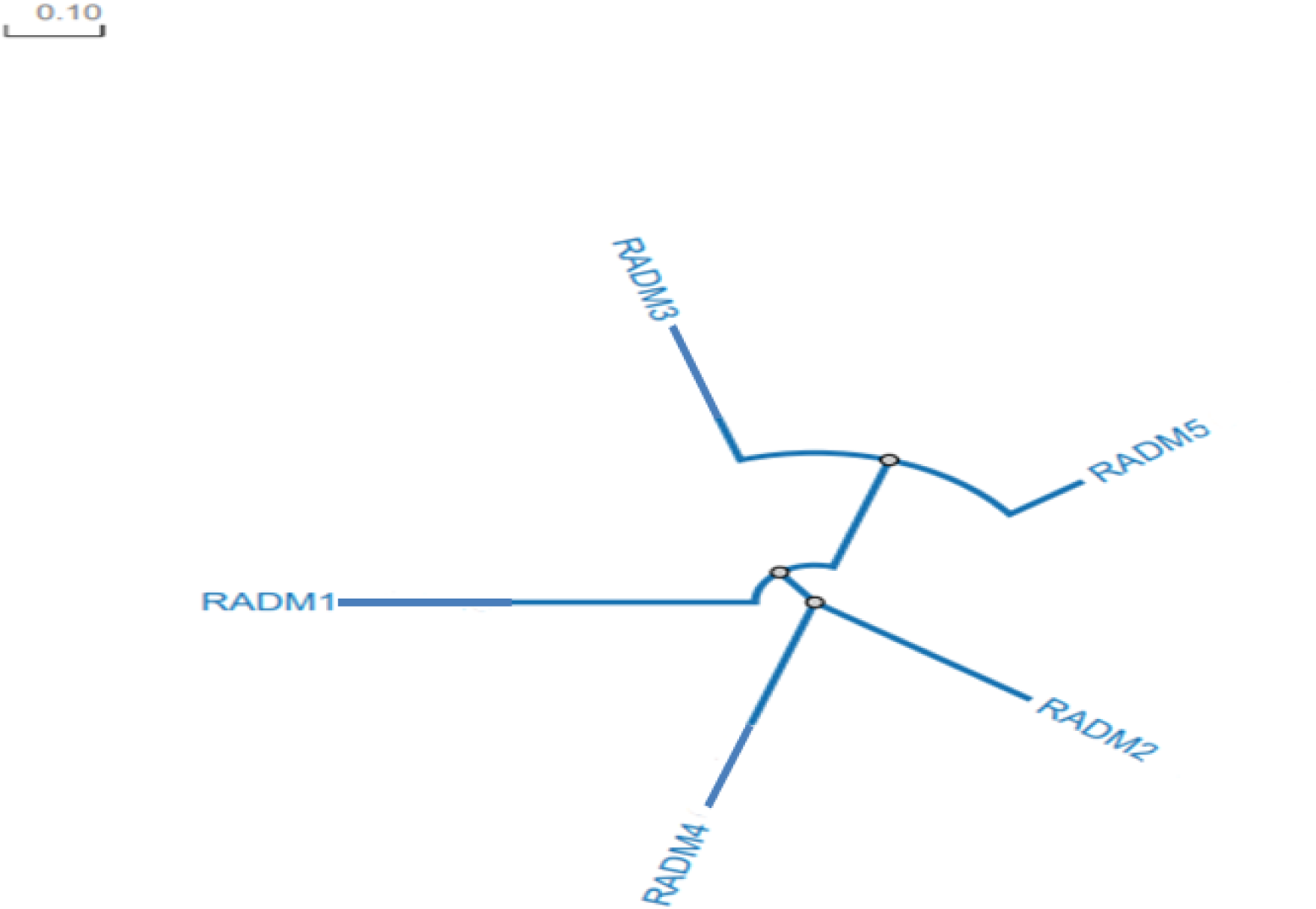
Phylogenetic grouping of actinobacterial isolate through Clustal Omega tool

**Table 13:** Molecular characterization of actinobacteria isolates.

| Isolate | 16S *r*RNA gene | Accession number | Family |
| --- | --- | --- | --- |
| RADM1 | *Streptomyces Clavuligerus* | PQ114139 | Streptomycetaceae |
| RADM2 | *Streptomyces Clavuligerus* | PQ120343 | Streptomycetaceae |
| RADM3 | *Rhodococcus erythropolis* | PQ114109 | Nocardiaceae |
| RADM4 | *Streptomyces Clavuligerus* | PQ120390 | Streptomycetaceae |
| RADM5 | *Rhodococcus erythropolis* | PQ039766 | Nocardiaceae |

### 2.6 Pot Experiment

On the basis of multiple and efficient PGP traits, stress resistance capabilities of isolated actinobacteria were further studied the effect of growth and drought tolerance on mung bean [*Vigna radiata* (L.)] plant using pot experiment in plant growth chamber. The result revealed that all actinobacterial isolates were able to stimulate the growth of mung bean seedlings under stress condition, which showed higher value of eco physiological parameters in comparison to control.

Actinobacteria, which generally represent an abundant proportion of soil microbiota, are particularly effective colonizers of plant root system and are capable of forming spores, they are able to endure unfavourable growth conditions and are more persistent in drought soils (Santos-Medellin *et al*., 2017). They play crucial role in organic matter cycling by increasing soil organic matter and nitrogen content along with essential macro and micro elements, which in turn ameliorate plant growth, carbon metabolism and allocation and improve plant yield (AbdElgawad *et al*., 2020).

During experiment seedlings were inoculated with bacterial isolates after 7 days and water application terminated after 14 days (T1 to T5) (Figure_13). After 21 days of sowing, in all the physiological parameters of mungbean seedlings treated with actinobacterial inoculats were studied in comparision to uninoculated seedlings (T6 and T7) at field capacity (Figure_13). All the bacterial treatments showed significantly better performance than the uninoculated drought induced treatment (T7) and uninoculated watered treatment (T6). Treatment T6 was continues with water supplementation for 21 days it showed significant growth with bacterial treatments. However, the treatment T7 in which application of water terminated after 14 days, revealed least growth for all the parameters than other treatments (Figure_14). Mungbean seedlings in T6 with continued water application were observed for growth parameters like, shoot length, root length, fresh weight and dry weight respectively 39.48%, 15.63%, 26.70%, and 19.80% higher than the drought induced treatment (T7).

**Figure 13:**
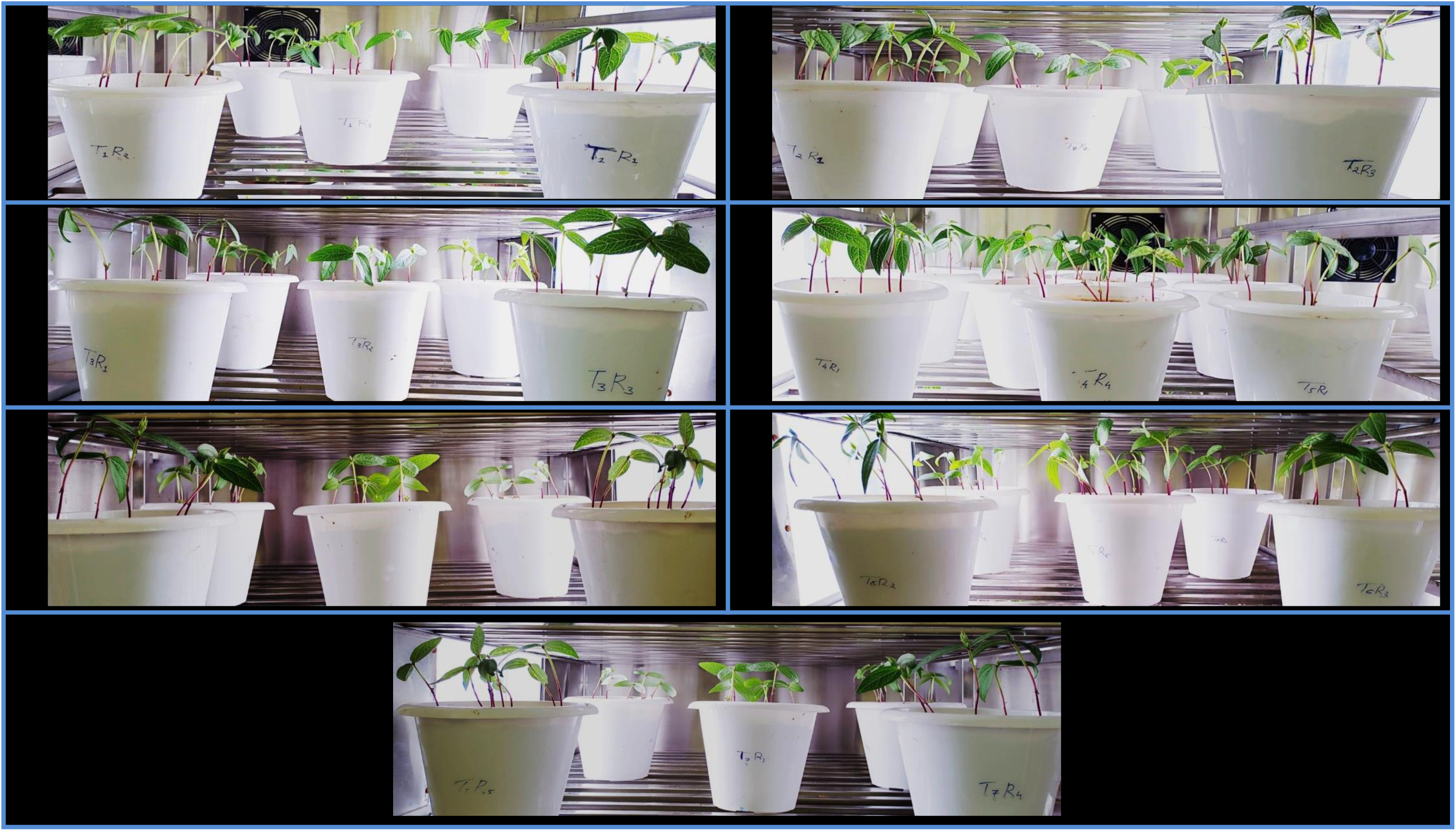
Mungbean seedling after seven days of bacterial inoculation and before drought induced

**Figure 14:**
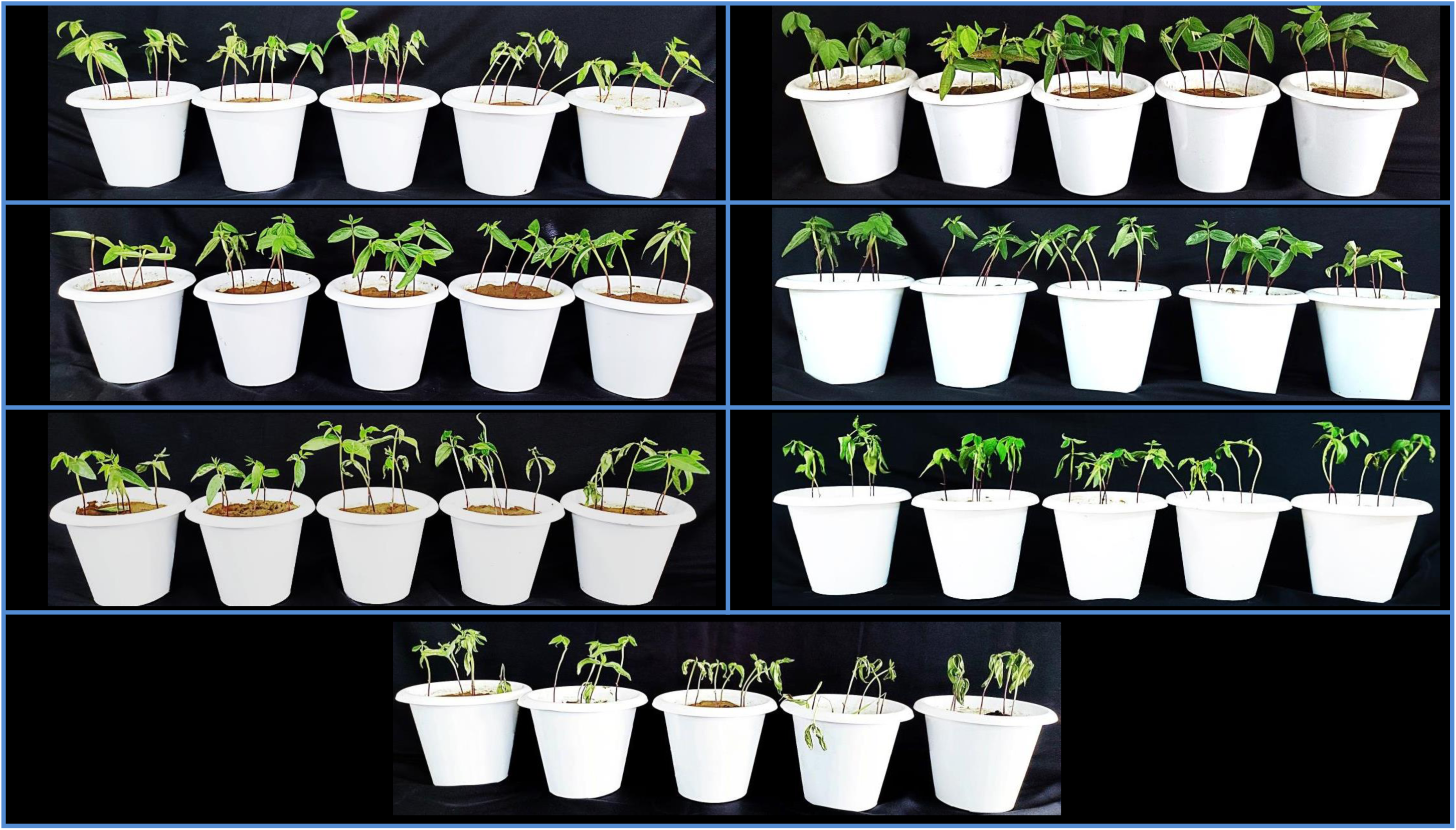
Mungbean seedlings after drought treatment and water resumes for one day

Mungbean seedlings inoculated with actinobacterial isolates showed variable growth parameters after 3 weeks (Figure_13). Significant increases in shoot length were observed in T2 (67.16% and 19.90%), T4 (61.31% and 15.70%), T1 (48.6% and 6.59%), T3 (44.38% and 3.53%), T5 (42.07% and 2.38%) as compared to T7 and T6 respectively. Significant increases in root length of seedlings were recorded in T2 (46.81% and 26.96%), T4 (44.57% and 25.03%), T3 (29.21% and 11.55%), T1 (29.03% and 11.58%), T5 (28.52% and 11.14%) as compared to T7 and T6, respectively. Increased fresh weights were observed in T2 (62.61% and 28.34%), T4 (58.35% and 24.98%), T3 (37.28% and 8.35%), T5 (35.76% and 7.15%), T1 (20.00% and 08.70%) as compared to T7 and T6, respectively. Similarly increased in dry weight of seedlings were recorded in T2 (60.69% and 34.12%), T4 (60.06% and 33.59%), T1 (37.40% and 14.68%), T3 (35.18% and 12.83%), T5 (32.96% and 10.97%) as compared to T7 and T6 respectively (Table 14).

**Table 14:**
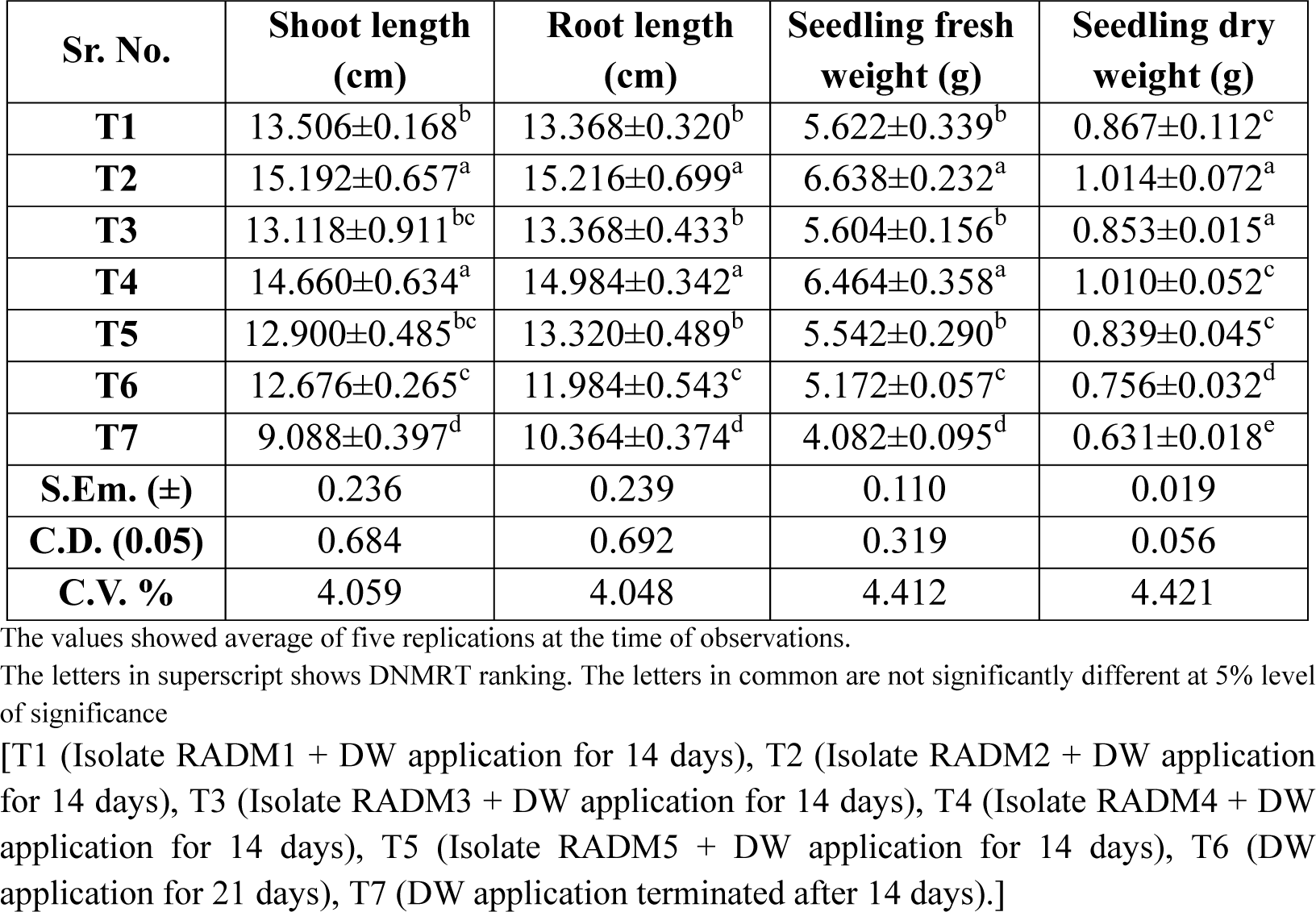
Growth improvement in mungbean by actinobacterial isolates under drought stress condition.

Changes in biochemical traits such proline, chlorophyll, and MDA (Malondialdehyde) content were observed under water stress in mungbean seedlings after 21 days. The water stress induced with actinobacterial treatments (T1 to T5) resulted in increased in proline, chlorophyll a, chlorophyll b and total chlorophyll contents and significant decrease in MDA content than treatments without bacterial inoculations (T6 and T7).

Accumulation of proline is common phenomenon that helps plants withstand drought stress and helps in acclimatization of crop plants which was examined and compared in mungbean. A profound effect of soil water crises was observed on proline content of seedlings (Table 15). Mungbean seedlings maintained low proline content in treatment T6 (1.91±0.278 µmol/g) at field capacity as compared with the drought induced treatment T7 (4.90±0.085 µmol/g). Significant increase of proline contents were found in actinobacterial treated seedlings than treatments without bacterial inoculation. Highest proline content was observed in the treatment T2 (6.41±0.095 µmol/g) followed by the treatment T4 (6.24±0.151 µmol/g), T1 (5.92±0.193 µmol/g), T3 (5.78±0.130 µmol/g), and T5 (5.57±0.110 µmol/g), respectively which revealed that the bacteria helped the plant to produce more proline content which protects the plant from drought stress through protein synthesis, scavaged reactive oxygen species, inhanced nutrient uptake, stabilized metabolism, chlorophyll synthesis, and root elongation.

**Table 15:**
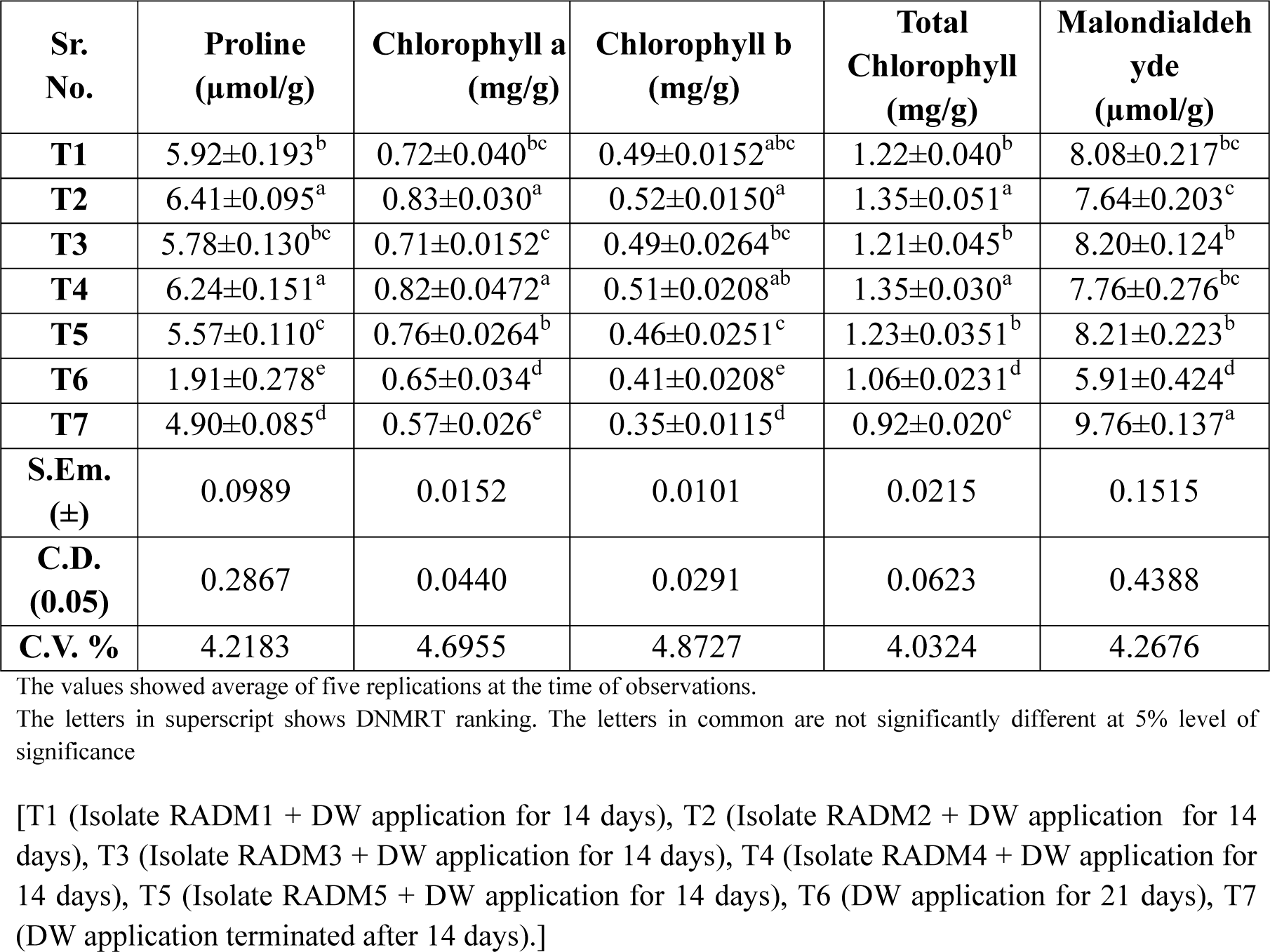
Oxidative stress features by actinobacterial isolates for drought tolerant in mungbean.

The plant pigment chlorophyll a, chlorophyll b and total chlorophyll content of seedlings were increased in bacterial treated seedlings under the influence of water stress (Table 15).

Observations revealed that significantly increased in chlorophyll a for the treatments T2 (45.61% and 27.69%), T4 (43.85% and 26.15%), T5 (33.33% and 16.92%), T1 (26.31 % and 10.76%), T3 (24.56% and 09.23%) as compared to T7 and T6, respectively. Chlorophyll b content were found significantly increased in bacterial treatment T2 (48.57% and 26.82%), T4 (45.71% and 24.39%), T3 (40.00% and 19.51%), T1 (40.00% and 19.51%) and T5 (31.42% and 12.19%) than the treatment T7 and T6, respectively without inoculation of actinobacteria. Similarly, increased in total chlorophyll content were recordedin leaves of seedling through bacteria inoculated treatment T2 (46.73% and 27.35%), T5 (33.69% and 16.03%), T4 (32.60% and 15.09%), T1 (32.60% and 15.09%) and T3 (31.52% and 14.15%) as compared to without inoculated treatments T7 and T6, respectively (Table 15).

Malondialdehyde (MDA) content in plant is a marker of oxidative stress and damage to plant membranes. The level of MDA in plants is used to evaluate the plant’s ability to tolerate stress. High level of MDA indicate damage to plant membranes and irreparable modification in protein and nucleic acids. Treatment without bacterial inoculation with drought induced T7 (9.76±0.137 µmol/g) recorded highest level of MDA content than all the other treatments. Plants with lower level of MDA under drought condition are generally considered more drought tolerant. Lowest MDA content was recorded in the treatment T6 (5.91±0.424) because of continued water supplement during experiment. Significant decrease in MDA content were recorded in Actinobacteria treated seedlings T2 (27.74%), T4 (25.77%), T1 (20.84%), T3 (19.02%) and T5 (18.87%) as compared to treatment without bacterial inoculated treatment T7 (Table 15).

## 3. Discussion

Sustainability is significant challenge currently being faced by human beings. How can we nourish the ever growing world population and at the same time offer viable soil for future crop production for next generation? (Glick, 2012). Plant growth promoting bacteria including rhizospheric bacteria and those which are free living in soil or associated to plant in rhizoplane, phyllosphere and inside the plant as endophytes. These microbes help plant growth by enhancing soil nutrient availability, the supply of phytohormones and provide systemic resistance induction against phytopathogens. Plant Growth promoting Bacteria (PGPBs) are a strong ally for sustainable agriculture. They offer an interesting alternative to chemical fertilizer and pesticides (Boukhatem at el. 2022).

Drought is one of the major environmental stresses that limit crop growth and productivity worldwide, while global warming and water scarcity will further make the situation worse. Thus, it is urgent to develop crop plants with improved drought tolerance. Several recent ecological studies have found that microbial symbionts can confer habitat specific stress tolerance to host plants, suggesting that the basis for the stress tolerance-enhancing effect of microbial symbionts (Alhuwalia et al. 2021).

Numerous microorganisms have been extensively studied for their plant growth-promoting rhizobacteria (PGPR) characteristics; however, actinobacterial species have gained attention primarily in the last twenty years for their beneficial properties. These bacteria are recognized for their significant metabolic capabilities, particularly in the production of a variety of secondary metabolites, including antibiotics. Additionally, actinobacteria contribute crucially to biogeochemical cycles and the processes involved in soil formation (Backer et al. 2018).

The phylum actinobacteria is a primary eubacterial phylogenetic clade containing diverse Gram’s positive bacteria that belong to several classes such as acidimicrobia, actinobacteria and thermoleophia. Actinobacteria are typically dominant soil microbe. They are important for the cycling of carbon, nitrogen, phosphorus, potassium and several other elements in soil. As saprophyte, they produce a range of extracellular hydrolytic enzymes which can degrade animal and plant polymers, including lignin, chitin, cellulose and other organic compounds.

Selection of rhizospheric soil of datura and khejri plant is because of a typical wild growing and desert plant species, is widely distributed in the arid, semi-arid and salinized regions of country. As we observed, there was no special treatment given or taken care for the growth, development and protection of these plants but still they are surviving in the environment, which attract the role of plant growth promoting bacteria in soil presented and their association with plant can be the one reason for survive of the plant against the abiotic stress conditions like drought, salinity, temperature or pH. The bacteria found in the rhizosphere have evolved multiple strategies to assist plants in managing the adverse effects of drought. These strategies may include alterations in phytohormones, which play a vital role in enabling plants to withstand environmental stresses, modifications to root structures, accumulation of osmolytes, enhancements in the plant’s antioxidant defense mechanisms, synthesis of exopolysaccharides, and the identification of particular genes that support plant growth and improve drought resilience.

Five morphologically different bacteria colonies of actinobacteria from the both the samples of datura and khejri’s rhizospheric soil were isolated in pure culture through serial dilution method. Isolates were marked sequentially 1 to 10 and stored in refrigerator at 4°C for further experimentation. The pure cultures were subsequently maintained on ISP2 in (International Streptomyces project 2) agar slants. These isolates underwent purification through repeated subculturing, followed by the isolation of single colonies to obtain pure cultures. The pure cultures were subsequently maintained on ISP2 agar slants according to the method outlined by Anjum and Chandra (2015).

All the ten actinobacterial isolates were screened based to their ability to survive and growth under drought condition *in-vitro*. For that all the isolates were tested in International Streptomyces project 1 (ISP 1) broth medium supplemented with the different concentration of polyethylene glycol (PEG8000) 5%, 10% and 15% of the and incubated for the different time intervals 24h, 48h and 72h and OD value were recorded. Comparison of tolerance for the growth of isolates at different concentration after different time of interval revealed that the least concentration of PEG8000 at 5% shows better growth than 10% and 15% concentration in medium. Isolate T9 had maximum tolerance to 5% PEG8000 (0.06627) and isolate T3 had least tolerance to 15% PEG8000 (0.127). So that, the five actinobacterial isolates which showed maximum tolerance against PEG8000 at 5% concentration according to the DNMRT value of comparing mean with different time interaction were selected and named as RADM1 [T9 (0.8473±0.0025 at 72h)], RADM2 [T8 (0.8473±0.0025 at 72h)], RADM3 [T10 (0.765±0.004 at 72h)], RADM4 [T4 (0.758±0.0017 at 72h)] and RADM5 [T2 (0.7453±0.0021at 72h)] for the further experiments.

Physiological characterization were done based on the ability to survive against their ability to grow with different temperature, pH and NaCl concentration ranges with media supplemented with 5% PEG8000. All the isolates had optimum growth at 40°C, among them isolate RADM4 has highest growth 0.91 ± 0.03 which were not significantly different from one another but significantly higher than the growth obtained for other isolates which followed by isolate RADM2 (0.89 ± 0.025) and the least growth was observed in isolate RADM5 (0.81 ± 0.025). At the highest temperature of 60°C, a significant decrease in the growth was observed for all the isolates.

The growth of bacteria was influenced by the pH levels in the medium. Bacterial isolates were identified within a pH range of 5 to 9, with all actinobacterial isolates exhibiting optimal growth at a pH of 7 compared to other pH levels. Among them isolate RADM2 (1.11± 0.05) and RADM4 (1.05± 0.05) had significantly higher growth compared to other isolates at pH 7 and the least growth was observed in isolate RADM3 (0.85± 0.04). Isolate RADM2 was able to withstand the at all the pH ranges than other isolates and at highest pH of 11 and lowest pH of 3 growth were observed 0.97± 0.025 and 0.29± 0.02 respectively which were significantly higher than the other isolates at that pH range.

All bacterial isolates were evaluated for their tolerance to salinity stress by cultivating them in media containing varying concentrations of sodium chloride (NaCl) at 0.5%, 1.0%, 2.0%, 3.0%, and 5.0%, along with an addition of 5% PEG. Significant differences were observed in the growth of bacterial isolates at different concentration. At 0.5% NaCl isolate RADM2 had highest growth 0.96± 0.04 followed by isolates RADM4 (0.95± 0.043) and RADM1 (0.88± 0.025) respectively.

Morphological evaluation plays a crucial role in the identification and characterization of microbial isolates through microscopic analysis. The isolates RADM1, RADM2, and RADM4 exhibited a rod-shaped morphology and were confirmed as positive for Gram staining, while RADM3 and RADM5 displayed variable forms, including both rod and coccus shapes, after a 48-hour incubation period. Additionally, the presence of filamentous mycelium, characterized by either branched or unbranched structures and clumps at the hyphal tips, was indicative of actinobacteria. On the agar plates, the actinobacteria colonies presented a yellowish brown coloration with a dry surface texture. All isolates exhibiting yellowish brown pigmentation on ISP2 culture media. The colonies displayed irregular forms and filamentous margins, characterized by indented peripheral edges (Anjum and Chandra, 2015).

Biochemical characterization includes carbohydrates utilization through IMVIC test kit revealed that all the bacterial isolates were found positive for indole, methyl red and glucose utilization and negative for voges proskauer’s and sorbitol utilization. Isolate RADM1 and RADM2 found positive for Citrate utilization arabinose, mannitol, sucrose utilization and negative for adonitol and lactose utilization. Isolate RADM3 were found positive for citrate, sucrose utilization, negative for mannitol, rhamnose utilization and produce intermediate reaction for adonitol, arbinose and lactose utilization. Isolate RADM4 were found positive for citrate, arbinose, mannitol utilization and negative for adonitol, lactose, rhamnose and sucrose utilization. Isolate RADM5 were found negative for adonitol, arabinose, lactose, mannitol, rhamnose and sucrose utilization.

All actinobacterial isolates demonstrated positive catalase activity during enzymatic characterization, as evidenced by the production of bubbles upon the addition of hydrogen peroxide, confirming their classification as Gram-positive bacteria. The catalase enzyme plays a crucial role in detoxifying harmful hydrogen peroxide, thereby protecting the bacteria. Additionally, the actinobacterial isolates exhibited positive results for nitrate reduction, indicated by the development of a cherry red colour following the addition of reagents A and B, which facilitates the availability of nitrogen sources essential for plant growth and development. However, the isolates were negative for the hydrolysis of starch into simpler sugar molecules, as no halo zone was observed in the *in vitro* tests.

The molecular characterization of bacteria is essential for the precise identification of bacterial species, elucidating their genetic relationships, monitoring the dissemination of specific strains within populations, and comprehencing their virulence factors, which is particularly significant across various domains of microbiology. The identification of actinobacterial isolates was conducted through 16S *r*RNA gene sequencing and subsequent analysis. The sequencing process utilized both forward and reverse primers. The resulting sequences, after alignment, were compared using the BlastN tool against the nucleotide sequences in the NCBI GenBank database with accession number PQ114139 (RADM1), PQ120343 (RADM2), PQ114109 (RADM3), PQ120390 (RADM4) and PQ039766 (RADM5). The 16S *r*RNA gene sequences from five isolates were submitted to NCBI GenBank, where accession numbers were assigned to the submitted sequences. Isolates RADM1, RADM2 and RADM4 were found respectively, 96.58%, 98% and 100% similarity to the actinobacterial species *Streptomyces clavuligerus*, which belongs to the order Streptomycetales within the phylum Actinomycetota. In contrast, isolates RADM3 and RADM5 demonstrated respectively 98% and 96% similarity to the species *Rhodococcus erythropolis*, classified under the order Mycobacteriales of the same phylum. The DNA sequences submitted to NCBI were utilized to construct a phylogenetic tree, incorporating bootstrap values to enhance the understanding of their evolutionary relationships based on genetic similarities and differences. The combined phylogenetic grouping of the actinobacterial isolates were prepared through Clustal Omega tool.

These two species of actinobacteria can used as promising alternative to chemical fertilizers and fungicides, serving as biofertilizers or bioinoculants. They facilitate plant growth through the production of various phytohormones and metabolites, enhance nutrient availability for plant growth and development, and offer protection against pathogens. These isolates have demonstrated their ability to improve physiological parameters, such as root and shoot length, as well as the fresh and dry weight of seedlings. Additionally, they contribute to the enhancement of oxidative stress responses, enabling plants to better withstand abiotic stress, including drought conditions.

Pot experiment include the inoculation of Actinobacterial isolates to the Mungbean seedlings and against the drought induced treatments with bacteria strains revealed the best pest performance for the physiological characterization and oxidative stress features in comparison with un inoculated continuous watered treatment and uninoculated drought induced treatment. Mung bean seedlings maintained seedlings growth parameters shoot length, root length, fresh weight and dry weight in treatment (T6- uninoculated watered treatment) 39.48%, 15.63%, 26.70%, and 19.80% higher than the drought induced treatment (T7- uninoculated drought induced treatment). Significant increases in shoot length were observed in T2 (67.16% and 19.90%), T4 (61.31% and 15.70%), T1 (48.6% and 6.59%), T3 (44.38% and 3.53%), T5 (42.07% and 2.38%) as compared to T7 and T6, respectively. Significant increases in root length of seedlings were recorded in T2 (46.81% and 26.96%), T4 (44.57% and 25.03%), T3 (29.21% and 11.55%), T1 (29.03% and11.58%), T5 (28.52% and 11.14%) as compared to T7 and T6 respectively. Increases fresh weights were observed in T2 (62.61% and 28.34%), T4 (58.35% and 24.98%), T3 (37.28% and 8.35%), T5 (35.76% and 7.15%), T1 (20.00% and 08.70%) as compared to T7 and T6 respectively. Similarly increases in dry weight of seedlings were recorded in T2 (60.69% and 34.12%), T4 (60.06% and 33.59%), T1 (37.40% and 14.68%), T3 (35.18% and 12.83%), T5 (32.96% and 10.97%) as compared to T7 and T6 respectively.

Drought conditions have led to alterations in the oxidative characteristics of plants, specifically in terms of proline levels, chlorophyll content, and malondialdehyde (MDA) concentrations. A significant impact of soil water deficiency was noted on the proline levels in seedlings. Mungbean seedlings exhibited a proline concentration of 1.91±0.278 µmol/g in treatment T7, which was maintained under field capacity, in contrast to the drought-affected treatment T6, which recorded a proline level of 4.90±0.085 µmol/g. Notably, seedlings treated with Actinobacteria demonstrated a marked increase in proline content compared to those without bacterial inoculation. The highest proline concentration was recorded in treatment T2 (6.41±0.095 µmol/g), followed by T4 (6.24±0.151 µmol/g), T1 (5.92±0.193 µmol/g), T3 (5.78±0.130 µmol/g), and T5 (5.57±0.110 µmol/g).

Increases were recorded in total chlorophyll content of seedling’s leaves of bacteria inoculated treatment T2 (46.73% and 27.35%), T5 (33.69% and 16.03%), T4 and T1 (32.60% and 15.09%), T3 (31.52% and 14.15%) as compared to without inoculated treatments T7 and T6 respectively. Chlorophyll is main component for the photosynthesis for the growth and development of plant. The presence of malondialdehyde (MDA) in plants serves as an indicator of oxidative stress and membrane damage. Elevated MDA levels suggest significant harm to plant membranes, as well as irreversible alterations in proteins and nucleic acids. Among the various treatments, the drought-induced T7 treatment, which did not involve bacterial inoculation, exhibited the highest MDA content, measuring 9.76±0.137 µmol/g, surpassing all other experimental conditions. Plants with lower level of MDA under drought condition are generally considered more droughts tolerant. Lowest MDA content was recorded in treatment T6 (5.91±0.424 µmol/g) because of continues water supplement during experiment. Significant decrease in MDA content were recorded in actinobacteria treated seedlings T2 (27.74%), T4 (25.77%), T1 (20.84%), T3 (19.02) and T5 (18.87%) as compared to treatment without bacterial inoculation T7.

## 4. Conclusion

Numerous microorganisms have been extensively studied for their plant growth-promoting rhizobacteria (PGPR) characteristics; however, actinobacterial species have gained attention primarily in the last twenty years for their beneficial properties. The Experiment revealed that the actinobacteria isolated from the rhizospheric soils have their own capability to survive under different environmental conditions. Bacteria were tested for their drought tolerance ability and survive against PEG8000. Selected five isolated were physiologically characterized through growing them with different temperature, pH and NaCl concentrations with drought induced treatment (with PEG8000) were found significant growth against abiotic stress condition. Morphological characteristics of bacterial isolates were found same as typical actinobacterial species like pigmentation, mycelial development and filamentous edges on culture medium. Biochemically all the isolates were able to utilize IMVIC and carbohydrate testes. All the actinobacterial isolates were found positive for nitrate reduction and catalase productions and negative for starch hydrolysis. Genomic studies of bacterial isolates revealed RADM1, RADM2 and RADM4 have maximum similarity with *Streptomyces Clavuligerus* and RADM3 and RADM5 had maximum similarity with *Rhodococcus erythropolis*. Inoculation of actinobacteria in pot experiment against the drought induced treatments of mungbean seedlings showed significant increase in physiological growth and oxidative stress condition of seedling against drought in comparison to uninoculated drought induced as well as continues water supply treatments. This showed that the use of actinobacteria as bioinoculant for drought tolerance and plant growth promotion can be batter alternative for natural farming and sustainable agricultural practices than chemical fertilizers as safe, protected and alive future.

## Acknowledgement

I acknowledge the Dean PG studies of Sardarkrushinagar Dantiwada Agricultural University for providing me all the facilities and equipments for my doctoral research.

